# Convergent energy state-dependent antagonistic signalling by CART and NPY modulates the plasticity of forebrain neurons to regulate feeding in zebrafish

**DOI:** 10.1101/2021.10.21.465223

**Authors:** Devika S. Bodas, Aditi Maduskar, Tarun Kaniganti, Debia Wakhloo, Akilandeswari Balasubramanian, Nishikant Subhedar, Aurnab Ghose

## Abstract

Dynamic re-configuration of circuit function subserves the flexibility of innate behaviours tuned to physiological states. Internal energy stores adaptively regulate feeding-associated behaviours by integrating opposing hunger and satiety signals at the level of neural circuits. Across vertebrate lineages, the neuropeptides CART and NPY have potent anorexic and orexic functions, respectively, and show energy state-dependent expression in interoceptive neurons. However, how the antagonistic activities of these peptides modulate circuit plasticity remains unclear.

Using behavioural, neuroanatomical and activity analysis in adult zebrafish, along with pharmacological interventions, we show that CART and NPY activities converge on a population of neurons in the dorsomedial telencephalon (Dm). While CART facilitates glutamatergic neurotransmission at the Dm, NPY dampens the response to glutamate. In energy-rich states, CART enhances NMDA receptor (NMDAR) function by PKA/PKC mediated phosphorylation of the NR1 subunit of the NMDAR complex. Conversely, starvation triggers NPY-mediated reduction in phosphorylated NR1 via calcineurin activation and inhibition of cAMP production leading to reduced responsiveness to glutamate.

Our data identify convergent integration of CART and NPY inputs by the Dm neurons to generate nutritional state-dependent circuit plasticity that is correlated with the behavioural switch induced by the opposing actions of satiety and hunger signals.

## INTRODUCTION

Adaptive regulation of feeding in response to internal energy needs is critical for survival. Internal sensory systems, located in the periphery and in the central nervous system, monitor energy homeostasis and signal satiety or hunger to the feeding-associated neural circuits. Hunger stimulates feeding circuits to generate motivational states that prioritise feeding and related activities over other behaviours. Conversely, satiety signals suppress the ‘feeding drive’ facilitating the pursuit of non-feeding-related behaviours. The flexibility of the feeding behaviour – exploiting or abandoning food resources – requires dynamic re-configuring of the activity states of the underlying circuits and the de-regulation of these processes forms the basis of eating disorders.

Neuropeptides, acting as hormones or local diffusive modulators convey nutritional state information to the nervous system (van den Pol, 2012). Neuropeptide-based neuromodulation of circuit function, via changes in neuron intrinsic properties or synaptic efficacy, may allow the same circuit to produce multiple outputs and behavioural outcomes (Nadim and Bucher, 2014). Neuropeptides typically engage G-protein coupled receptors (GPCRs) which, via intracellular signalling, may change circuit properties and extend acute signals of internal needs into long-lasting state changes mediated by biochemical hysteresis. For example, in the rodent arcuate nucleus (Arc), orexigenic neuropeptide ghrelin initiates an AMPK-mediated positive-feedback to potentiate glutamate release for extended periods (Yang et al., 2011). Consequently, several studies have implicated neuropeptides in mediating transitions between distinct behavioural states (Root et al., 2011; Elbaz et al., 2012; Flavell et al., 2013; Betley and Sternson, 2015).

Cocaine- and amphetamine-regulated transcript (CART) and Neuropeptide Y (NPY) are abundantly expressed neuropeptides conserved across vertebrate lineages. Both the neuropeptides have been implicated in a diverse range of physiological functions, including energy homoeostasis and food intake (Lau and Herzog, 2014; Subhedar et al., 2014; Singh et al., 2021)

CART is a potent anorexigenic agent and considered to be an endogenous satiety factor (Kristensen et al., 1998), especially in the framework of the Arc in the hypothalamus (Singh et al., 2021). Expression of CART increases in response to feeding in the Arc (Kristensen et al., 1998) and recent chemogenetic studies underscore the requirement of CART peptide in Arc neurons to inhibit food intake (Farzi et al., 2018). In contrast, hypothalamic NPY is orexigenic, and promotes food intake and energy conservation (Zhang et al., 2019). Starvation increases NPY expression in the Arc (Sainsbury and Zhang, 2010) and NPY-signalling from AgRP-expressing Arc neurons is critical for the sustained maintenance of the feeding drive (Chen et al., 2016).

Within the Arc, the coordinated co-regulation of the AgRP/NPY and the POMC/CART containing neurons in response to signals indicating metabolic states is well documented (Dietrich and Horvath, 2013; Andermann and Lowell, 2017). The physiologically opposing activities of these two populations and their effect on “second-order” neurons that drive satiety is a major regulator of the feeding behaviour (Garfield et al., 2015; Andermann and Lowell, 2017). However, the intracellular signalling engaged by these neuropeptides in the “second-order” neurons to re-configure the circuit properties and drive homoeostatic plasticity remains inadequately defined.

In zebrafish, NPY and CART are expressed in several brain areas, including the periventricular hypothalamus (homologous to mammalian Arc; Forlano and Cone, 2007) and the entopenduncular nucleus (EN) (Mathieu et al., 2002; Mukherjee et al., 2012; Yokobori et al., 2012; Akash et al., 2014). Starvation decreases CART mRNA expression in the periventricular hypothalamus and the EN (Nishio et al., 2012; Akash et al., 2014) while increasing NPY expression (Yokobori et al., 2012).

Treatment with glucose also increases CART protein expression in the EN (Mukherjee et al., 2012). The zebrafish periventricular hypothalamus and the EN appear to be nutritional state sensitive interoceptive areas regulating energy homeostasis, in part via CART/NPY. Accordingly, treatment with NPY increases feeding in zebrafish and its action is mediated by the NPY Y1 receptor (Yokobori et al., 2012).

Despite the prominent involvement of CART and NPY in vertebrate energy homeostasis, it remains unclear how their activities re-configure the downstream, “second-order” circuits to meet behavioural demands. In this study, using zebrafish, we show that CART and NPY shape the feeding drive to match the prevailing energy states. The opposing energy state information represented by the two neuropeptides are integrated by a group of forebrain neurons by transforming the antagonistic signalling into dynamic changes in glutamatergic neurotransmission.

## MATERIALS AND METHODS

### Animals

Adult, wild-type zebrafish (*Danio rerio*) of both sexes were housed in stand-alone housing systems and maintained on 14hL/10hD cycle. Fish were fed Ziegler feed and live artemia 3 times a day. For calcium activity imaging experiments, adult Tg(NeuroD:GcAMP6f) in the *nacre* background were used. All behavioural assays and activity imaging experiments were performed between 10 am and 5 pm. No randomization methods were used to allocate animals to different experimental groups. The Institutional Animal Ethics Committee approved all the procedures employed in this study. This study was not pre-registered.

### Chemicals

The following reagents were administered to the zebrafish brain via intracerebroventricular (icv) delivery. The concentrations indicated reflect the amount delivered by the icv route. Rat CART peptide (55-102) (2 pmol in 1x PBS, Phoenix, Cat. No. 003-62), full length human NPY peptide (10 pmol in 1x PBS, Sigma-Aldrich, Cat. No. N5017), anti-CART antibody (1:500; gift from Drs. Lars Thim and Jes Clausen, Novo Nordisk, Denmark); D -(+)-glucose monohydrate (40 nmol in 1x PBS, glucose; Sigma-Aldrich, Cat. No. 49161), BIBP-3226 (100 pmol in 0.1% DMSO, Sigma-Aldrich, Cat. No. B174), MK801 (0.015 nmol in 1x PBS, Sigma-Aldrich, Cat. No. 77086), AP5 (0.1 nmol in 1x PBS, Tocris, Cat. No. 0106), KT5720 (1.4 pmol in 1x PBS, PKAi; Sigma-Aldrich, Cat. No. K3761), GF109206X (1.5 pmol in 1x PBS, PKCi; Tocris, Cat. No. 0741), FK506 (0.13 nmol in 1x PBS, Sigma-Aldrich, Cat. No. F4679), Forskolin (1.2 nmol in 0.1 % ethanol, Sigma-Aldrich, Cat. No. F6886). The following reagents were used for procedures not requiring icv administration. 2-Phenoxy Ethanol (1:2000 in system water, Sigma-Aldrich, Cat. No. P1126), D -(+)-glucose monohydrate (Sigma-Aldrich, Cat. No. 49161), sucrose (Sigma-Aldrich, Cat. No. S9703), paraformaldehyde (Sigma-Aldrich, Cat. No. 158127), poly-L-lysine (Sigma-Aldrich, Cat. No. P8920), normal goat serum (Sigma-Aldrich, Cat. No. 9023), bovine serum albumin (BSA) (SRL Pvt. Ltd., Cat. No. 83803), 1, 1’, dioctadecyl-3,3,3’, 3’-tetramethylindo-carbocyanineperchlorate (DiI; Invitrogen Cat. No. D3911).

### Intracerebroventricular (icv) injections

Fish were anaesthetised in 2-phenoxy ethanol (1:2000). The anaesthetised fish were placed on a cotton bed containing anaesthesia solution to submerge its head. Icv delivery protocol was modified after Yokogawa et al., 2007. Briefly, pharmacological agent or vehicle was delivered directly into the ventricular space using an 31G needle (Becton Dickinson Insulin Syringe U40 - 31G) attached via a catheter to a 10 μl Hamilton microsyringe. After injection, the fish were returned to their tanks and allowed to recover before proceeding for either behavioural recordings or immunohistochemistry. Unless otherwise indicated, 30 mins following injection, the animals were subjected to behavioural task or sacrificed and the tissues processed for immunofluorescence studies.

### Immunofluorescence

Fish were subjected to icv injection with drug or vehicle and allowed to recover for 30 mins. After recovery, fish were anesthetised and craniotomised to expose the dorsal surface of the brain. Samples were fixed in 4% PFA overnight at 4 °C. After fixation for 12-14hrs (10 hrs for pNR1 staining), brain was dissected and cryoprotected with 25% sucrose prior to sectioning. Serial 15-20 µm thick sections of the entire telencephalon were mounted onto lysine coated slides and stored at −40 °C till further processing. Sections were allowed to dry for 2 hrs and washed with 0.5% Triton X-100 in 1x PBS (PBST) three times for 10 min each). PBS of the following composition was used: NaCl, 137 mM; KCL, 2.7 mM; Na_2_HPO_4,_ 10 mM; KH_2_PO_4_, 1.8 mM; pH-7.2. Sections were then blocked using 5% BSA in PBST for 1 hr prior to the addition of primary antibodies and incubated overnight at 4°C. Next day, the washes were repeated and sections blocked in 5% BSA/ 5% HIGS for 1 hr. The following primary antibodies were used in this study: anti-p-ERK (1:700, Cell Signalling technologies, Cat. No. 9101), anti-p-ERK (1:500, Abcam, Cat. No. ab50011), anti-pNR1(Ser-897) (1:70, Millipore, Cat. No. ABN99), anti-Synaptophysin (1:100, Invitrogen, Cat. No. PA1-1043).

Secondary antibodies were added and incubated at room temperature (RT) for 2.5 hrs in dark. The following secondary antibodies were used: Anti-rabbit Alexa Flour 488/568 (1:500, Invitrogen, Cat. No. A-11034/ A-11036) and anti-mouse Alexa Flour 488/568 (1:500, Invitrogen, Cat. No. A-11029/A-11031). Sections were washed and mounted in media (0.5% N propyl gallate, 70% glycerol, 1M tris pH8.0) containing 4′,6-diamidino-2-phenylindole (DAPI) (Invitrogen, Cat. No. D1306).

Sections were observed under Axioimager Z1 (Zeiss) epifluorescence microscope and representative images were acquired using confocal microscope (SP8, Leica or LSM 780, Zeiss). In order to ensure reliable comparisons across different groups and maintain stringency in tissue preparation and staining conditions, all the brain sections were processed concurrently under identical conditions.

### Feeding Behaviour

Fish were isolated from home tanks and housed singly in the experimental tank prior to behavioural analysis. Fig. S1 outlines the protocol followed for all feeding behaviour experiments. Briefly, for first two days of habituation, the experimental tanks were moved to the recording chamber and the fish were fed floating, insoluble food pellets (15 ± 5 pellets; Taiyo) for 1 hour before returning to the housing rack. For habituation to injection and handling stress, the fish were anesthetised and delivered mock icv injection (saline, 0.9% NaCl) on day 4 and day 5. Fish were allowed to recover in recording chamber for 1 hour, before returning the tanks to the housing racks. Depending on the experiment, the fish were either starved or given pellet food for 1 hr on these days. Fish were observed on all days of the protocol for any anxious/abnormal behaviour. Fish exhibiting such behaviours were not included for further analysis. On the day of the experiment, fish were anaesthetized and icv injected with appropriate reagents or vehicle. Following a recovery period of 15 mins, floating pellets of food (~15 ± 5) were added to the tank and the behaviour of the fish recorded individually using a video recorder (Sony Handycam DCR SR-47 and DCR SR-20) for an hour. All recordings were conducted under conditions of uniform lighting and temperature. In all experiments, animals deprived of food for 2.5 days are referred to as ‘starved’ and those that received food as per regular feeding schedule, and used for experiments within one hour of feeding, are referred to as ‘fed’. The recorded videos were not blinded and the number of biting attempts was counted manually. The data was plotted as the cumulative number of biting attempts displayed during 60 mins.

### Calcium imaging

Starved, adult Tg(NeuroD:GcAMP6f) fish were injected with either vehicle or drugs. After 15 min, the fish were anesthetized in ice cold HEPES-based Ringer’s solution [NaCl, 134 mM; MgCl_2_, 1.2 mM; CaCl_2_, 2.1 mM; KCl, 2.9 mM; HEPES, 10 mM; Sucrose, 10 mM (for glucose-deprivation) or Glucose, 10 mM (to mimic satiety); pH 7.2] bubbled with 100% O_2_ and craniotomised to dissect the telencephalon. The optic nerves were cut off and the telencephalon was dissected out by cutting just rostral to habenula. The telencephalon was carefully lifted from the base while removing the ventral connections and immediately transferred to Ringer’s solution supplemented with 100% O_2_.

For *ex vivo* whole-brain preparations, starved fish were anesthetised in ice-cold HEPES-based Ringer’s solution. First the optic nerves and connections on the ventral side were severed and the whole brain was dissected out by separating it from spinal cord.

The intact telencephalon or the whole brain was mounted in 4% low-melt agarose (in 50% Ringer’s) on to a RC-26 GLP chamber from Warner Instruments (Cat. No. 640236). The imaging chamber was mounted onto an upright confocal microscope (Leica TCS SP8 MP) and continuously perfused with HEPES-based Ringer’s solution via am VC 8 perfusion system (Warner Instruments, Cat. No.640186) directly triggered by the recording software (Leica). Images were acquired at 9.8 Hz at a resolution of 512 x 300 pixels using a 488 nm laser in the resonant, bi-directional scanning mode and a 25x water-immersion objective (N.A 0.95). Baseline imaging was performed for 1 min before the start of the glutamate stimulation. Glutamate (75 µM, in Ringer’s buffer) was perfused from the 2^nd^ to 5^th^ mins and subsequently washed out.

### Data Analysis

Immunofluorescence: The tissue slices were observed under Axioimager Z1 (Zeiss, 20x objective 0.8 NA) with the imaging conditions kept constant across all treatment groups. The number of p-ERK positive cells in Dm were scored manually. To avoid over estimation of cell count due to sectioning, the cell numbers were corrected using Abercrombie’s method (Abercrombie, 1946).

Average pNR1 intensity per cell in Dm was calculated using IMAGE J. The exposure time and other imaging conditions were kept constant for every sample processed. Each neuron in Dm was selected using manual thresholding on the DAPI filter, and the mean intensity of pNR1 per neuron was calculated using IMAGEJ in-built algorithms. Images were acquired using a 40x objective (1.3 NA) on a SP8 confocal microscope (Leica).

Calcium imaging: Measurements of population level calcium activity in Dm were performed using ImageJ and the average GCaMP6f fluorescence in the Dm at each time point was calculated. For this, we manually marked an ROI encompassing the Dm using the wand tool in ImageJ on maximum intensity projection of all the frames. This ROI was then applied to the time lapse series and the baseline fluorescence (F0) estimated by taking the average of fluorescence values of 620 frames before the start of glutamate treatment. The relative change in fluorescence (ΔF/F0), was calculated using the formula F-F0/F0, where F stands for the fluorescence value at a given time. The maximum response amplitude, i.e., maximum ΔF/F0 was determined from the temporal sequence using Microsoft Excel. The total magnitude of the response was calculated by computing the area-under-the-curve (AUC), from start of glutamate treatment to end of the washout period, in the GraphPad Prism ver 8 software.

Feeding assay: Videos were analysed manually at 1.5x speed and the number of biting attempts were scored. The data were distributed in bins of 15 mins and cumulative biting attempts were plotted using GraphPad Prism ver 8 software.

### Statistical Analysis

Data were tested for normality using the Shapiro-Wilk test in GraphPad Prism 8. Behaviour and immunohistochemical data were subjected to the t-test with Welch’s correction or the Mann-Whitney test (for single comparison) and One-Way ANOVA with the Tukey’s test or Two-way ANOVA, with Bonferroni’s post-hoc analysis (for multiple comparisons). All values are expressed as mean ± SEM of the group in the Results section and differences were considered significant at *p*<0.05. Graphs were plotted using the GraphPad Prism 8.0 software. Data for feeding drive are represented as columns (with error bars marking mean± SEM) of cumulative biting attempts in 15 min bins over a 1-hour period. Change in GCaMP6f fluorescence over time is represented as XY plots with the dark line representing mean and the dotted lines denote the SEM. All data shown as violin plots have the thicker dashed line representing median and dotted lines representing first and third quartile.

## RESULTS

### CART neuropeptide mediates satiety during energy-rich metabolic states

To quantitatively evaluate the feeding drive of adult zebrafish, we developed a behavioural assay that scores the number of biting attempts made by the animal upon presentation of solid, floating food pellets (see Methods and Fig. S1). We found that hungry fish made many more biting attempts over a period of 1 h (794±57.46; Fig. 1 A) compared to fed animals (338.8±70.4; Fig. 1 A). Analysis of the number of biting attempts across 15 min bins revealed cumulated difference over the 1 h period. To rule out contributions from altered locomotion or stress-induced behaviours, we evaluated animals under fed and starved conditions for locomotion kinematics and indicators of anxiety-associated behaviours but found no change in these parameters (Fig. S2).

**Figure 1.**
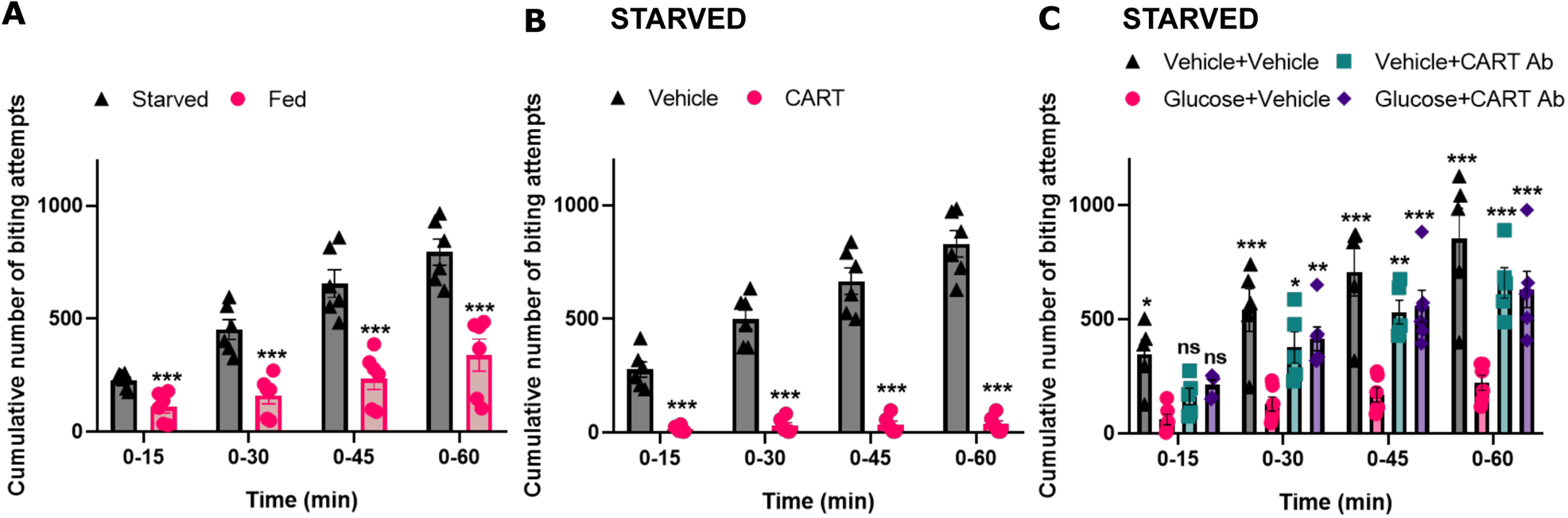
CART neuropeptide regulates satiety in adult zebrafish. **(A)** The feeding drive as indicated by the cumulative number of biting attempts made by fed and starved animals in 15 min bins (N=6 animals per group). **(B)** The cumulative number of biting attempts made by starved fish intracerberoventicularly injected with either vehicle or CART peptide (N=6 animals per group). **(C)** The cumulative number of biting attempts made by starved fish icv injected with either vehicle, glucose, CART antibodies or coinjected with immunoneutralising CART antibodies (CART Ab) along with glucose (N= 5 for vehicle control and CART Ab treated animals and N=6 for glucose and CART Ab + glucose treated animals). Data are represented as cumulative biting attempts in 15 min bins over a 1-hour period and compared using two-way ANOVA, with Bonferroni’s post-hoc analysis for significance in comparison with the glucose treated starved fish (error bars represent ± SEM; ns, not significant; ***p<0.001, **p<0.01).

To directly assess the role of CART neuropeptide in anorexia, we delivered CART peptide (55-102 aa) via the intracerebroventricular (icv) route to the brain of starved fish. Strikingly, exogenous administration of CART directly to the brain was sufficient to induce acute anorexia in starved fish (36.21±14.89; Fig. 1 B) compared to starved animals injected with vehicle (829.4±58.03 Fig. 1 B). However, CART treatment did not induce anxiety-like behaviours or modify locomotion (Fig. S2).

To test if central application of glucose resulted in anorexia, glucose was directly introduced via the icv route to the brains of starved fish to mimic energy surfeit conditions and preclude peripheral inputs. In contrast to starved animals (852±133.3), glucose-injected fish displayed a dramatic reduction in biting attempts (221.2±33.67; Fig. 1 C). As CART promotes anorexia (Lau and Herzog, 2014b) and CART peptide expression is sensitive to metabolic states in the zebrafish brains (Mukherjee et al., 2012; Akash et al., 2014), we used an immunoneutralising antibody against the CART peptide to test if endogenous CART signalling mediates the glucose-induced reduction in feeding. Indeed, glucose-induced anorexia in starved animals was alleviated in the presence of CART immunoneutralising antibodies also introduced by the icv route (629.9±79.31; Fig 1 C). CART antibodies on their own did not influence the feeding behaviour (658.4±67.13; Fig 1 C).

Collectively, these results establish the role of CART neuropeptide signalling in the zebrafish brain in mediating the reduction in food intake behaviour during energy-rich metabolic states.

### CART signalling alters the activity of telencephalic Dm neurons

The entopeduncular nucleus (EN) in the ventral telencephalon as well as periventricular hypothalamic nuclei, like the nucleus lateralis tuberis (NLT), show energy-state dependent changes in CART mRNA expression (Akash et al., 2014). Administration of glucose also increased the number of CART peptide expressing cells in the EN and the NLT (Fig. S3; Mukherjee et al., 2012).

CART immunofluorescence analysis of transverse sections of the zebrafish telencephalon was used to evaluate the projections from the EN. CART-positive EN soma was found to project to the dorsomedial telencephalon (Dm; Fig S4 A). This is consistent with earlier reports indicating that EN neurons project to the Dm (Turner et al., 2016). We used the lipophilic dye, DiI, to confirm if the Dm receives projections from EN and other energy status-responsive interoceptive areas. DiI-positive fibres from the Dm typically extended caudally and ventrally and merged with the lateral forebrain bundle. Labelled cells were found in the ipsilateral EN (Fig S4 B,C). Fibres were also traced caudally into the hypothalamus and labelled somata were located in the preglomerular nucleus and the NLT (Fig S4 B,D). This connectivity is consistent with previous reports from teleost fish, including zebrafish (Yáñez et al., 2017). Dual immunostaining for CART peptide and the presynaptic marker synaptophysin revealed significant co-localisation, indicating the presence of CART in presynaptic terminals at the Dm (Fig S4 F). However, no CART expressing somata were detected in the Dm.

Collectively, these results demonstrate that CART peptide expression in EN and the periventricular hypothalamus is controlled by metabolic states and these areas send projections to the Dm (Fig. S4 E). This connectivity suggested that Dm neurons could be responsive to energy state-dependent CART activity. To assess the involvement of Dm, we used phosphorylated ERK (p-ERK), a marker of recent neuronal activity (Randlett et al., 2015).

Starved fish injected with CART peptide via the icv route had an elevated number of p-ERK positive neurons in the Dm (533.9±89.04) in comparison to starved, vehicle-injected animals (159.5±34.52) (Fig 2 A-C). As these data suggest a CART-mediated increase in neuronal activity in the Dm, we tested if feeding was sufficient to induce neuronal activation of the Dm neurons by comparing starved and recently fed fish (where endogenous levels of CART are high; Fig S3). Remarkably, Dm also shows increased neuronal activity (423.2±26.61) in fed fish as compared to starved controls (180.0±12.42) (Fig 2 D-F). The subpopulation of Dm neurons responding to CART/fed states was limited to a ~ 250 µm region rostral to the level of the anterior commissure (Fig 2 G-I).

**Figure 2.**
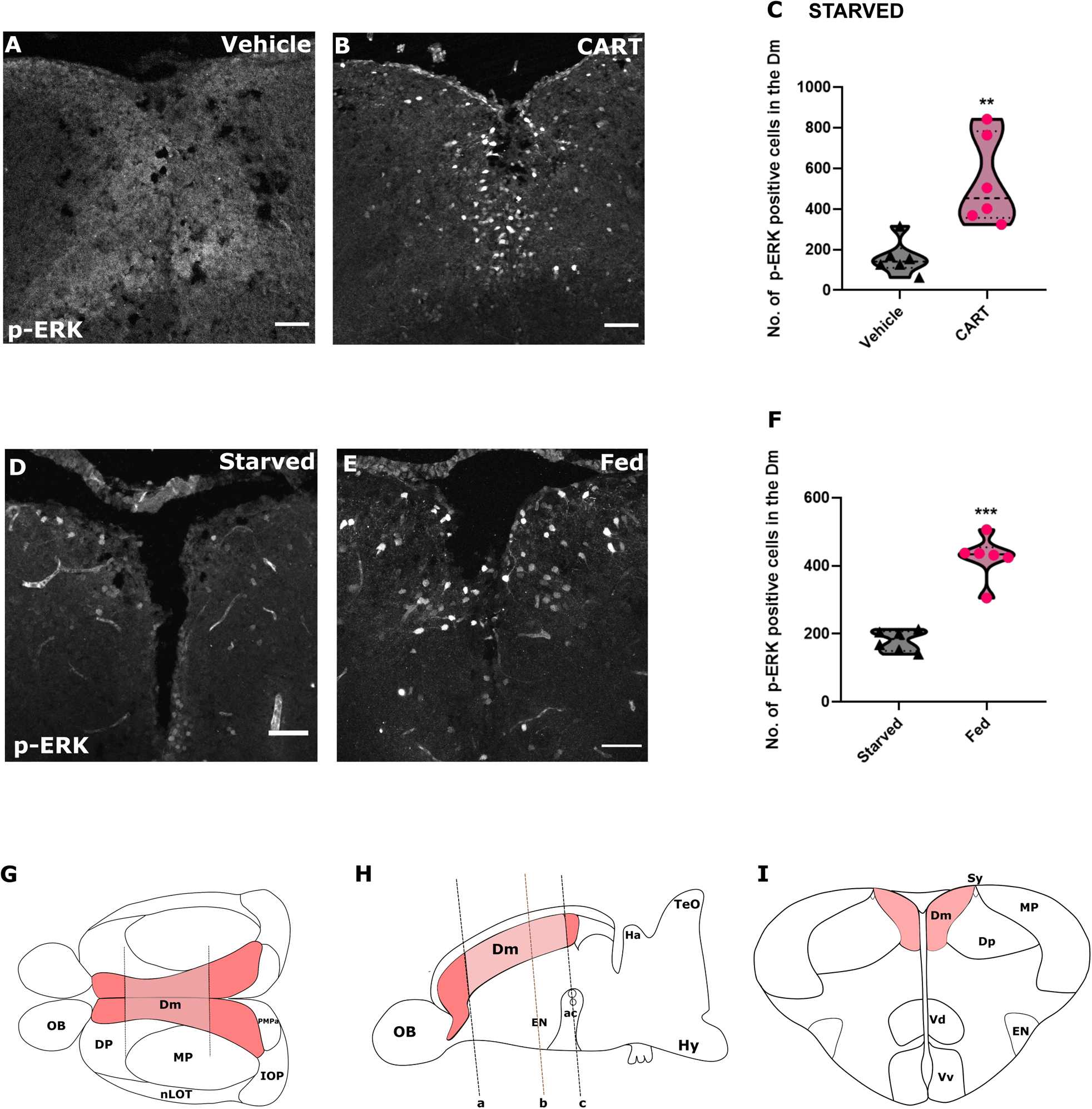
CART neuropeptide and satiety increases the activity of the dorsomedial telencephalon (Dm) neurons. **(A,B)** Representative micrographs of Dm region from the transverse sections of the telencephalon showing p-ERK immunoreactive (p-ERK-ir) cells in starved fish icv injected with either vehicle or CART peptide. **(C)** The number of p-ERK-ir cells in Dm in starved fish icv injected with either vehicle or CART. The data are compared using the unpaired t-test with Welch’s correction (N=6 animals per group; **p<0.01). **(D,E)** representative micrographs of Dm region from the transverse sections of the telencephalon showing p-ERK-ir cells under starved and fed conditions. **(F)** The number of p-ERK-ir cells in Dm under starved and fed states. The data are compared using the unpaired t-test with Welch’s correction (N=6 animals per group). (**p<0.01, ***p<0.001). Scale bar: 50 µm. **(G)** The top view of zebrafish telencephalon with shaded region (dark red) indicating the entire dorsomedial telencephalon. The area bounded by dotted lines within dorsomedial telencephalon (light red) shows differential activity upon CART treatment and under fed/starved conditions. We refer to this subregion as Dm for the purposes of this study. **(H)** Side view of zebrafish brain with dark red area marking entire dorsomedial telencephalon. The region of our interest – within dorsomedial telencephalon - Dm (light red) is marked by dotted lines (a: marks the rostral limit of the region which shows differential activity; b: marks the plane of section shown in I; c: marks the caudal limit at the level of the anterior commissure (ac); the distance from a to c is ~250 µm). **(I)** A transverse section of telencephalon with the Dm region marked in light red colour. [ac, anterior commissure; Dp, posterior zone of dorsal telencephalon; EN, entopeduncular nucleus; Hy, hypothalamus; Ha, habenula; IOP, integrative olfactory pallium; MP, medial pallium; nLOT-nucleus of the lateral olfactory tract; OB, olfactory bulb; PMPa, posteromedial pallial nucleus; Sy, sulcus yipsiloniformis; TeO, optic tectum; Vd, dorsal nucleus of ventral telencephalic area; Vv, ventral nucleus of ventral telencephalic area]

These findings suggest that enhancement of CART function at the Dm, possibly by the energy state responsive neurons of the EN and the periventricular hypothalamus, results in increased neuronal activity. However, the possibility of additional inputs from other CART-ergic sources cannot be ruled out

### CART modulates NMDAR signalling to regulate the feeding behaviour and the activity of Dm neurons

The dorsal telencephalon of teleosts, including the Dm, is rich in glutamatergic neurons (Aoki et al., 2013) and shows significant binding of labelled kainate and l-glutamate (Tong et al., 1992) implicating glutamatergic neurotransmission. Studies in rodents have previously implicated NMDA receptor (NMDAR)-mediated CART signalling in the context of innate fear processing (Rale et al., 2017) and nociceptive transmission (Chiu et al., 2010) in rat. We investigated NMDAR signalling in CART-mediated regulation of feeding behaviour using NMDAR antagonists. Icv delivery of AP5 (competitive NMDAR antagonist) alone in starved fish showed no difference in biting attempts (845.3±89.82; Fig 3 A) as compared to vehicle controls (812.8±84.38. However, CART-induced anorexia (47.33±11.4) was abrogated when CART peptide and AP5 were co-administered (488.7±42.03; Fig 3 A). Application of the non-competitive NMDAR antagonist, MK801 produced similar results (Fig S5 A).

**Figure 3.**
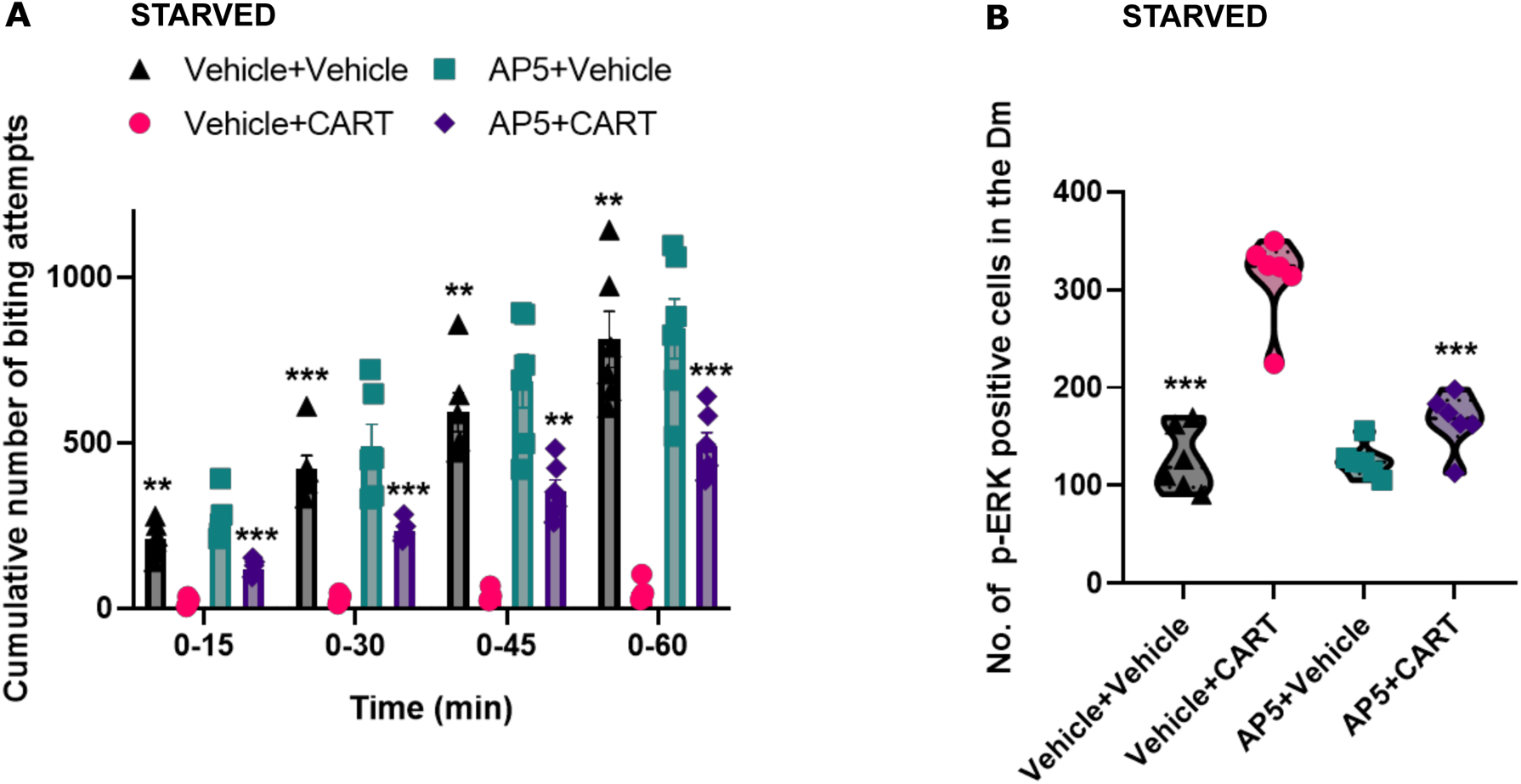
NMDAR signalling is necessary for CART-induced anorexia and increased activation of Dm neurons. **(A)** The cumulative number of biting attempts made by starved zebrafish icv injected with either vehicle, CART peptide, AP5 or coinjected with AP5 and CART peptide. (N=6 animals per group). Data are represented as cumulative biting attempts in 15 min bins over a 1-hour period and compared using two-way ANOVA, with Bonferroni’s post-hoc analysis (error bars represent ±SEM; **p<0.01, ***p<0.001). **(B)** The number of p-ERK-ir cells in starved fish icv injected with either vehicle, CART, AP5 or coinjection of AP5 along with CART. The data are compared using two-way ANOVA, with Bonferroni’s post-hoc analysis for significance in comparison with the CART treated starved fish (N=6 animals per group; ***p<0.001).

AP5 was co-injected with CART peptide in starved animals to evaluate the role of NMDAR signalling in modulating the activity of the Dm neurons. Neuronal activity in the Dm, indicated by the number of p-ERK positive neurons, was robustly attenuated in CART peptide and AP5-treated fish (165.7±11.87) compared to animals injected with only CART peptide (312±18.2; Fig 3 B). AP5 alone (125.2±6.96) did not influence the number of p-ERK positive Dm neurons as compared to those in vehicle controls (126.7±13.35; Fig 3 B). The application of MK801 produced similar results (Fig S5 B).

Together these data indicate that NMDAR activity is required for processing CART-induced satiety signals in the Dm neurons.

### PKC and PKA phosphorylate NR1 to mediate the modulation of NMDAR signalling by CART

Protein kinase C (PKC) and Protein kinase A (PKA) mediated phosphorylation of the NR1 subunit of the mammalian NMDAR is known to potentiate NMDAR signalling by increasing the channel opening probability and enhancing calcium permeability (Lan et al., 2001; Skeberdis et al., 2006). PKA and PKC have also been implicated in CART-signalling in *in vitro* experiments (Chiu et al., 2010). We next evaluated PKA and PKC function in CART signalling using specific pharmacological inhibitors of PKA and PKC kinase activity.

KT5720 (selective PKA inhibitor, PKAi), when co-injected into the ventricle of starved fish along with CART peptide, alleviated (697.49±105.59) the strong reduction in the number of biting attempts induced by CART peptide alone (30.33±10.61 Fig. 4 A). The inhibitor on its own had no significant effect on the feeding drive (713.92±215.68) as compared to starved control animals (782.9± 71.70; Fig. 4 A).

**Figure 4.**
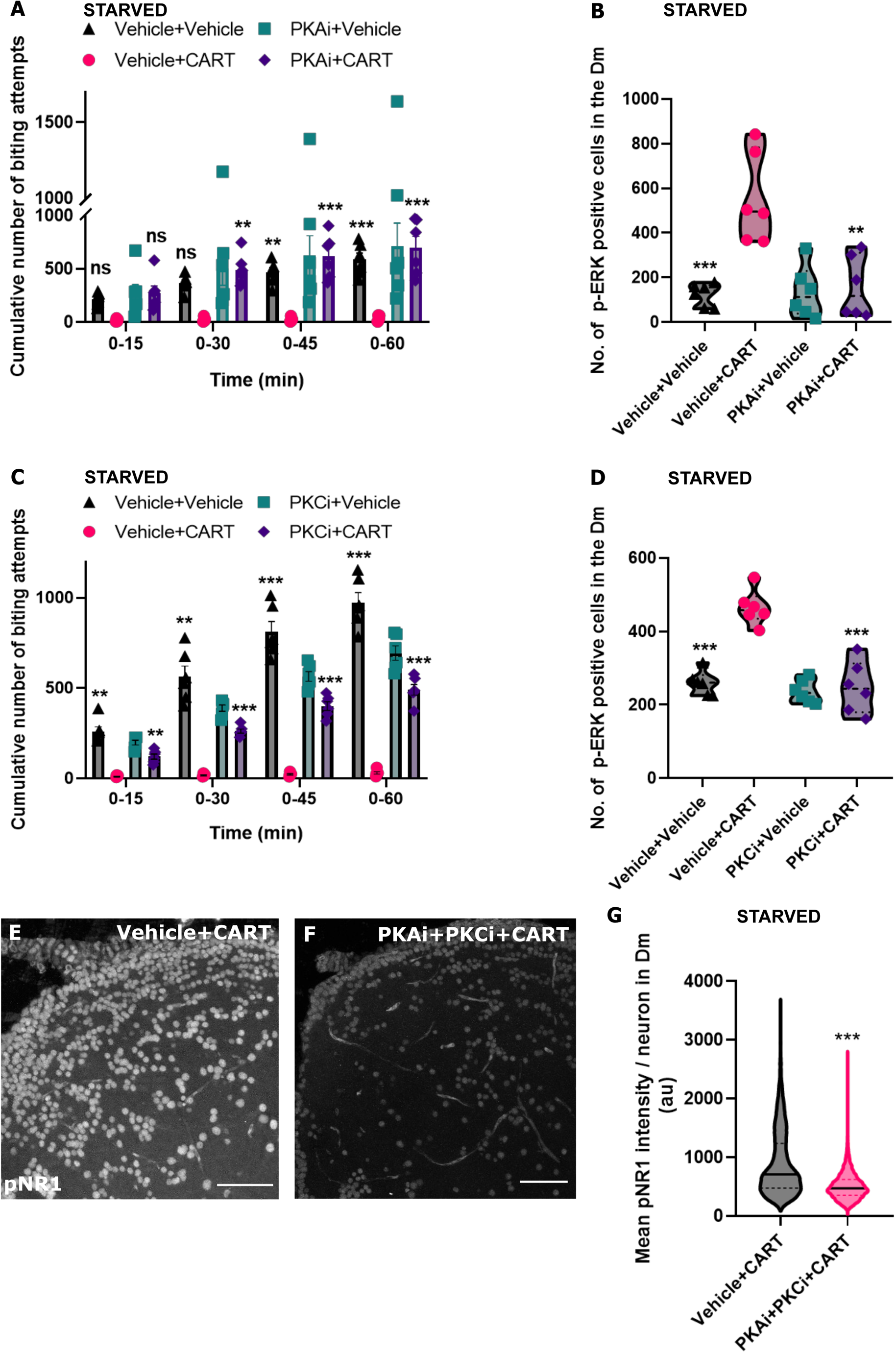
CART-induced anorexia and activation of Dm neurons require PKA and PKC activity. **(A)** The cumulative number of biting attempts made by starved fish icv injected with either vehicle, CART, PKAi or coinjection of PKAi along with CART. (N=6 animals per group). Data are represented as cumulative biting attempts in 15 min bins over a 1-hour period and compared using two-way ANOVA, with Bonferroni’s post-hoc analysis for significance in comparison with the CART treated starved fish (error bars represent ±SEM; ns, not significant; ***p<0.001, **p<0.01). **(B)** The number of p-ERK-ir cells in starved fish icv injected with either vehicle, CART peptide, PKAi or coinjected with PKAi and CART peptide. Data are compared using two-way ANOVA, with Bonferroni’s post-hoc analysis for significance in comparison with the CART treated starved fish (N=6 animals per group; ***p<0.001, **p<0.01). **(C)** The cumulative number of biting attempts made by starved fish icv injected with either vehicle, CART peptide, PKCi or coinjected with PKCi and CART peptide. (N=6 animals per group). Data are represented as cumulative biting attempts in 15 min bins over a 1-hour period and compared using two-way ANOVA, with Bonferroni’s post-hoc analysis for significance in comparison with the CART treated starved fish (error bars represent ±SEM; ***p<0.001, **p<0.01). **(D)** The number of p-ERK-ir cells in starved fish icv injected with either vehicle, CART, PKCi or coinjection of PKCi along with CART. The data are compared using two-way ANOVA, with Bonferroni’s post-hoc analysis for significance in comparison with the CART treated starved fish (N=6; *** p< 0.001). **(E,F)** representative micrographs of one lobe of the Dm region from the transverse sections of the telencephalon showing phosphoNR1 (Ser-897) (pNR1) immunoreactivity in starved fish icv injected with either CART peptide or coinjected with PKAi and PKCi along with CART peptide. **(G)** Quantification of mean pNR1 intensity per neuron (au: arbitrary units) in the Dm of starved fish icv injected with either vehicle and CART or coinjected with PKAi and PKCi along with CART peptideThe data were compared using unpaired t-test with Welch’s correction (12253 cells in vehicle +CART group and 12015 cells in PKAi + PKCi + CART group; N=3 animals per group; ***p<0.001. Scale bar: 50 µm.)

Parallel studies evaluating the activity of Dm neurons revealed a robust suppression of CART-induced neuronal activation in fish co-administered with PKAi and CART peptide (158.1±56.47; Fig. 4 B) compared to those injected with only CART peptide (554.5±82.75; Fig. 4 B).

As observed for PKAi, inhibition of PKC by GF109206X (selective PKC inhibitor, PKCi) also reversed the suppression of the feeding drive by CART peptide (489.8±29.38 for PKCi + CART and 29.83±6.76 for CART alone; Fig. 4 C) and abrogated the activation of Dm neurons by CART (247.7±28.99 for PKCi + CART and 464.9± 19.57 for CART alone; Fig. 4 D).

Together, these data implicate PKA and PKC activity in CART-mediated activation of Dm neurons and satiety. PKC and PKA are known to phosphorylate mouse NR1 at multiple residues, including serine 897 (Chen and Roche, 2007). Specific antibodies against phosphoserine 897 of the mouse NR1 (pNR1) were used to evaluate the status of NR1 phosphorylation in the Dm neurons. While the Dm of CART-treated animals displayed strong pNR1 fluorescence (892.1±5), co-injection of PKAi, PKCi, and CART peptide greatly reduced the pNR1 signal (505.9±2.02) (Fig. 4 E-G).

Collectively, these data identify CART signalling mediated phosphorylation of NR1 via PKA and PKC. The heightened neuronal activity of Dm neurons could be attributed to the potentiation of NMDAR function driven by NR1 phosphorylation.

### CART treatment enhances NMDA receptor function resulting in increased sensitivity of Dm neurons to excitatory stimuli

Phosphorylation of NR1 subunit of NMDARs by PKA and PKC is known to enhance NMDAR signalling (Lan et al., 2001; Skeberdis et al., 2006). We speculated that a CART-induced increase in phosphorylation of NMDARs would sensitise and enhance the response of Dm neurons to glutamatergic inputs. We tested this hypothesis directly using the Tg(NeuroD:GCaMP6f) zebrafish, which shows strong expression of the genetically-encoded calcium sensor, GCaMP6f in the Dm. *Ex vivo* preparations of the intact telencephalon were dissected from starved animals treated earlier with either saline or CART via the icv route (see Methods and Fig. 5 A). Dm neurons of vehicle-treated animals failed to respond to glutamate stimulation.

**Figure 5.**
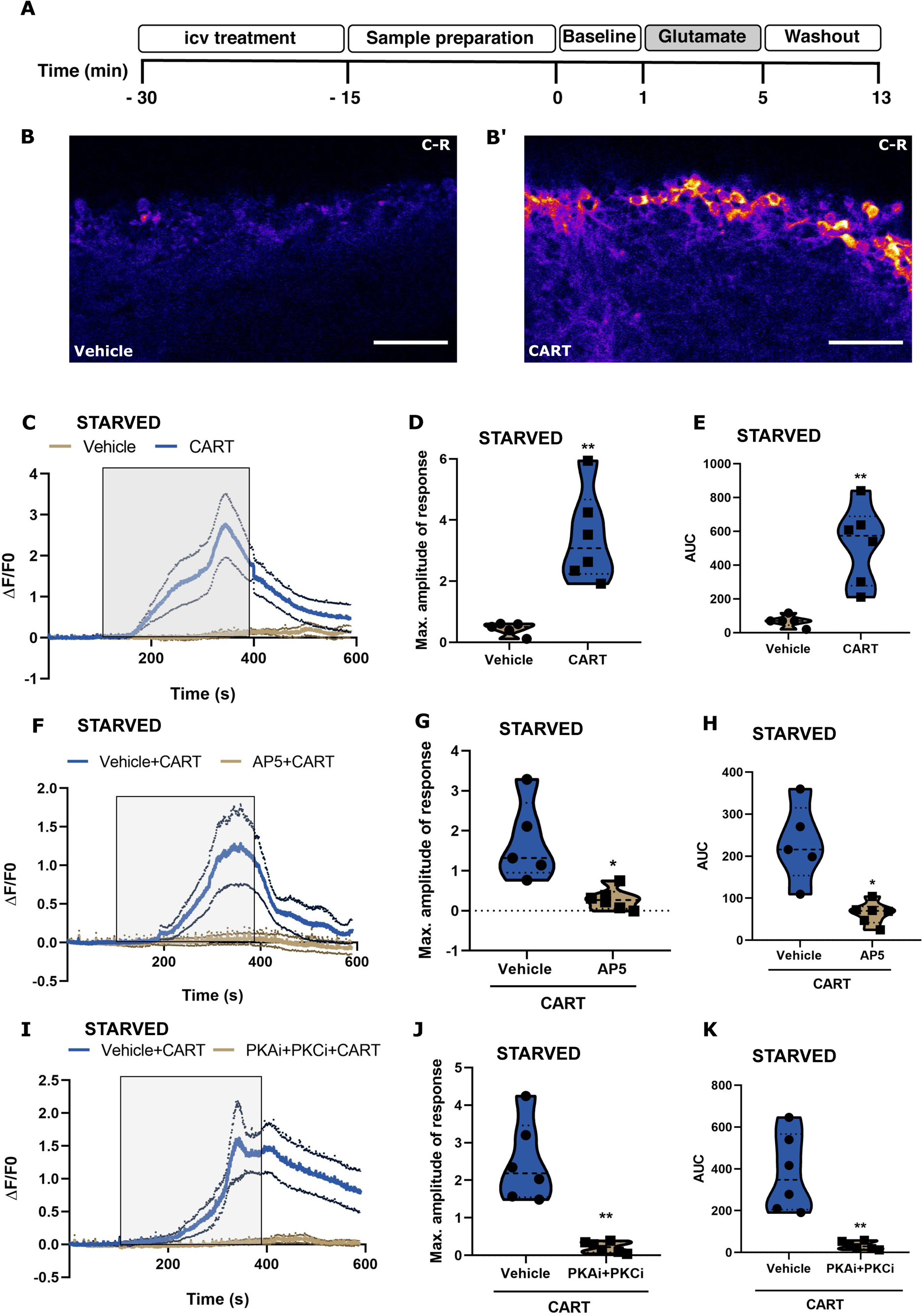
CART treatment enhances NMDAR function resulting in increased sensitivity of Dm neurons to glutamate. **(A)** A timeline of the experimental protocol followed for all calcium imaging experiments using *ex-vivo* telencephalon preparations (details in Methods). Representative micrographs of neural activity (GCaMP6f fluorescence) in the Dm of vehicle (**B**) and CART peptide (**B’**) injected <u>Tg(NeuroD:GCaMP6f) transgenic</u> <u>animals</u> following stimulation with glutamate for the same time. The ventricle is towards the top of the image and C and R indicate caudal and rostral directions, respectively. Scale bar: 50 µm. In all graphs **(C, F, I)**, the traces represent relative change in fluorescence intensity (ΔF/F0) across time. The dark line represents the mean and the dotted lines denote the standard error of mean. The shaded rectangular box indicates the duration of glutamate presentation. (**C**) The change in fluorescence intensity (ΔF/F0) of Dm neurons in starved fish treated with either CART peptide (blue trace) or vehicle (brown trace). **(D)** The maximum response (ΔF/F0) and **(E)** the extent of the total response – AUC (area under the curve) for Dm neurons in starved fish treated with either CART peptide or vehicle. The data were compared the t-test with Welch’s correction. (N=6 telencephalons per group, **p<0.01). **(F)** The change in fluorescence intensity (ΔF/F0) of Dm neurons in starved fish treated with either CART peptide (blue trace) or coinjected with AP5 and CART peptide (brown trace). **(G)** The maximum response (ΔF/F0) and **(H)** the extent of the total response of Dm neurons in starved fish treated with either CART peptide or coinjected with AP5 and CART. The data were compared using t-test with Welch’s correction (N= 5 for CART and N=6 for AP5+CART telencephalons, *p<0.05). **(I)** The change in fluorescence intensity (ΔF/F0) of Dm neurons in starved fish treated with either CART peptide (blue trace) or coinjected with PKAi and PKCi along with CART peptide (brown trace). **(J)** The maximum response (ΔF/F0) and **(K)** the extent of the total response of Dm neurons in starved fish treated with either CART peptide or coinjected with PKAi, PKCi along with CART peptide. The data were compared using using t-test with Welch’s correction (N=6 telencephalons per group, **p<0.01).

However, CART peptide-treated animals showed heightened activity of Dm neurons in response to glutamate. Both the maximum amplitude of the response (0.44±0.09 for vehicle control and 3.43±0.6 for CART) and the magnitude (area under the curve) of the response (69.04±15.29 for vehicle control and 522.6±94.66 for CART) was substantially higher in starved animals receiving CART peptide (Fig. 5 B-E). Co-treatment of AP5 along with CART peptide prevented the activation of the Dm neurons by glutamate (maximum amplitude:0.29±0.11 compared to 1.724±0.44 for CART alone; magnitude: 64.76±10.88 compared to 230.7± 41.33 for CART alone) and underscored the involvement of NMDAR signalling (Fig. 5 F-H).

To determine if the change in the glutamate sensitivity of Dm neurons in response to CART required PKA and PKC activity, we co-administered selective PKA and PKC inhibitors along with CART. Interestingly, blocking PKA and PKC activity abolished the CART-induced increased sensitivity of Dm neurons to glutamate (Fig.5 I-K). The maximum amplitude of the response was reduced to 0.21±0.06 (PKAi+PKCi+CART) from 2.48±0.43 (CART alone), and the magnitude of the response dropped from 380.3±75.86 (CART alone) to 33.39±7.71 (PKAi+PKCi+CART).

Together these data suggest that CART signalling via PKA and PKC sensitizes NMDA receptors in the Dm neurons, possibly via post-translational modification of NR1 subunits, leading to the enhanced activation of these neurons. This CART neuropeptide-mediated response of the Dm neurons may constitute a neural representation of the sated state of the animal.

### The activity of Dm neurons is tuned to changes in energy states

To assess if Dm exhibits heightened activity in energy-rich metabolic states, we used *ex vivo* whole-brain preparations from starved animals maintained in sucrose (to maintain gluco-deprived conditions). Following initial evaluation of the response of Dm neurons to glutamate stimulation, the preparation was incubated in glucose to mimic energy-rich conditions and re-evaluated for neural activity in response to glutamate stimulation (see Methods and Fig. 6 A).

**Figure 6.**
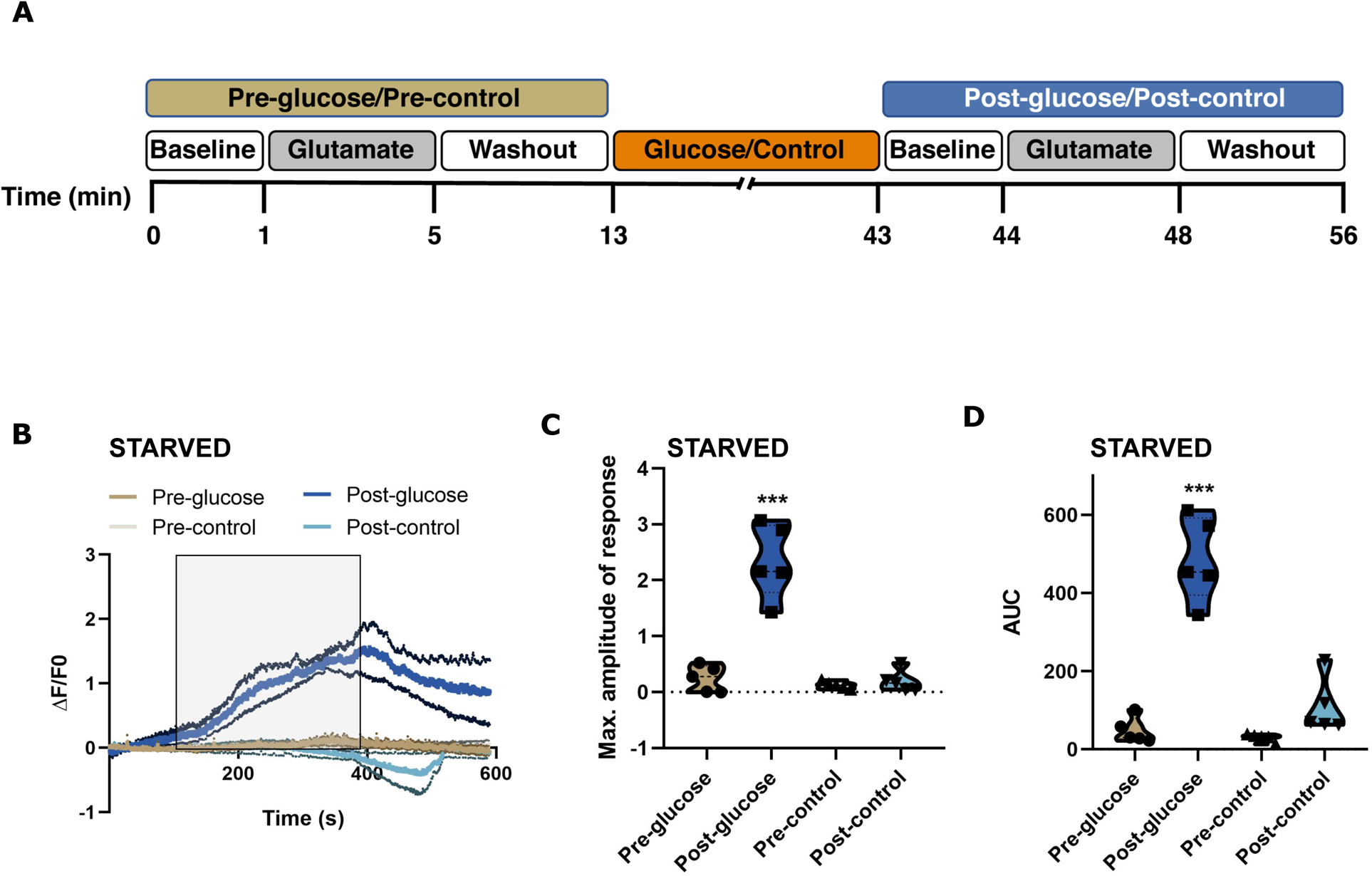
Activity of Dm neurons increases in response to glucose. **(A)** A timeline of the experimental protocol followed for calcium imaging experiments using *ex-vivo* whole brain preparations (details in Methods) **(B)** The traces represent relative change in fluorescence intensity (ΔF/F0) over time in response to glutamate. The thicker line represents mean while the associated finer lines indicate ± SEM. The shaded rectangular box indicates the duration of glutamate presentation. Response of Dm neurons in *ex-vivo* whole brain samples post 30’ incubation with glucose (dark blue trace; post-glucose) is substantial compared to pre-glucose treatment (dark brown trace). Control experiments showed no response of Dm neurons before (light brown; pre-control) and after (light blue; post-control) 30 min incubation in glucose-free media. **(C)** The maximum response (ΔF/F0) and **(D)** the extent of the total response of Dm neurons pre- and post-incubation with glucose/glucose-free media. The data were analysed using one-way ANOVA and Tukey’s multiple comparisons test (N=5 brains in each group, ***p<0.001).

Glutamate stimulation of whole-brains of starved animals maintained in gluco-deprived conditions (pre-glucose) failed to activate Dm neurons (maximum amplitude of response: 0.243±0.1 and magnitude of response: 48.29±14.71; Fig 6 B-D).

However, upon incubation of the same whole-brain preparation with 10 mM glucose for 30 mins (post-glucose), the Dm neurons responded strongly to glutamate stimulation (maximum amplitude of response: 2.33±0.29 and magnitude of response: 485±48.05; Fig 6 B-D). In control experiments, whole-brain preparations from starved animals were maintained throughout in glucose-free media, including the 30 min period between the two glutamate stimulations. In these control experiments, Dm neurons did not respond to glutamatergic stimulation either before (pre-control) or after the 30 min control incubation (post-control; Fig 6 B-D).

These experiments indicate that activity in the Dm is tuned to changes in glucose levels in the brain and neuronal activity in Dm is correlated to energy states. As CART levels are sensitive to metabolic states (Fig S3; Mukherjee et al., 2012; Akash et al., 2014), the glucosensitive neurons of the EN and hypothalamus may contribute to elevating endogenous CART signalling at the Dm neurons to mediate this effect. Interestingly, the change in Dm activity appears to be independent of peripheral mediators of energy homeostasis.

### NPY promotes food intake in zebrafish

Beyond sensing energy sufficiency, interoceptive awareness is also expected to involve signals indicating depletion of energy stores. Neuropeptide Y (NPY) is a potent orexigenic neuropeptide (Zhang et al., 2019), which is expressed in the EN and periventricular hypothalamic regions of teleosts, including zebrafish (Mathieu et al., 2002; Mukherjee et al., 2012; Akash et al., 2014; Turner et al., 2016) and regulates the feeding behaviour (Yokobori et al., 2012). We tested if NPY signalling regulated zebrafish feeding drive in a manner obverse to that of CART.

A selective NPY receptor Y1R (Y1R) antagonist, BIBP-3226 (BIBP; Doods et al., 1995; Yokobori et al., 2012; Kaniganti et al., 2021), was administered to starved fish to evaluate endogenous NPY signalling in regulating the feeding drive. Compared to vehicle-injected starved animals (789±69.46), BIBP introduced by the icv route robustly suppressed the biting frequency (144.7±21.57; Fig. 7 A) and is consistent with earlier reports (Yokobori et al., 2012).

**Figure 7.**
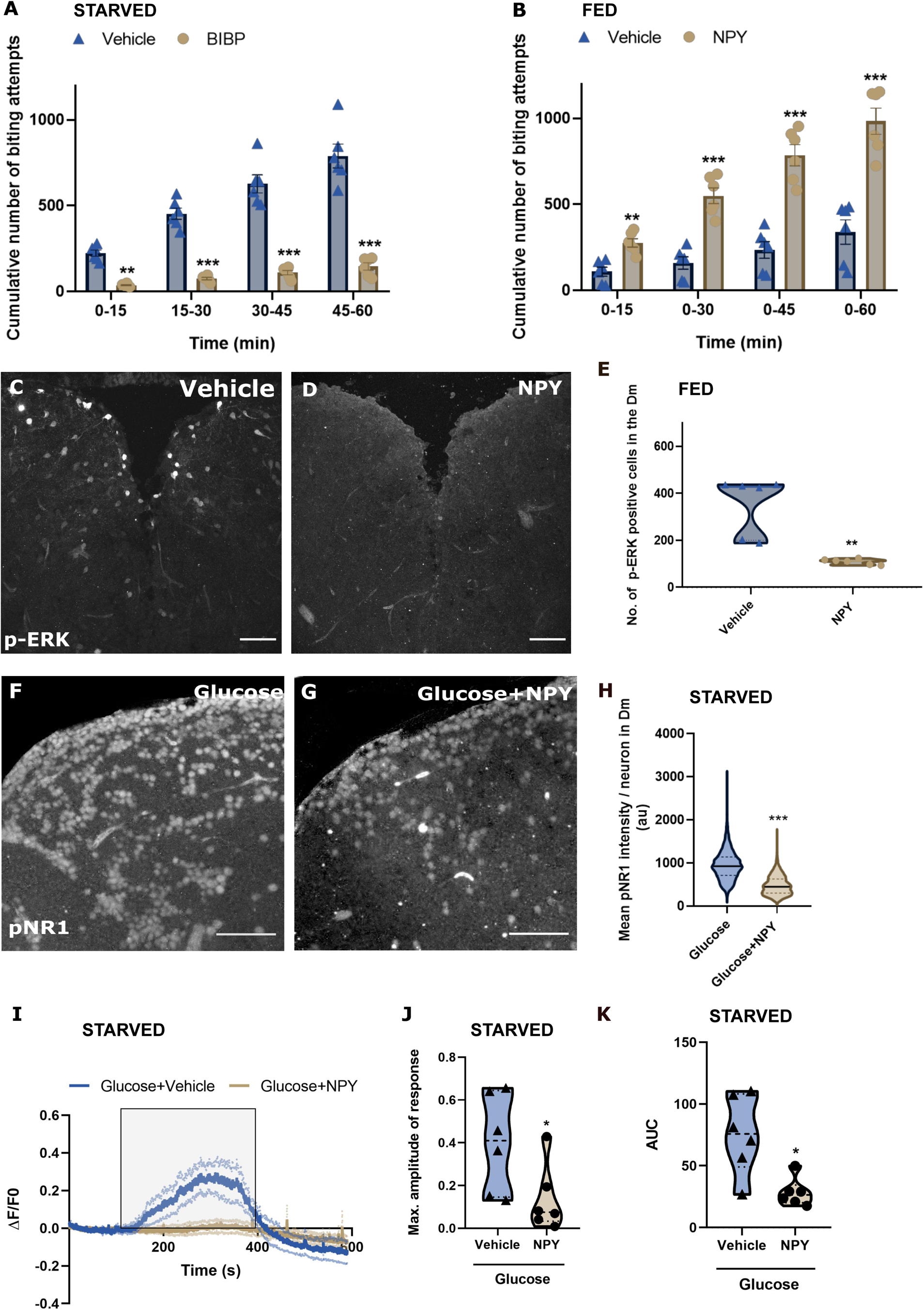
NPY promotes food intake and reduces the activation of Dm neurons. **(A)** The cumulative number of biting attempts made by starved fish icv injected with either vehicle or BIBP3226 (N= 6; ***p<0.001). **(B)** The cumulative number of biting attempts made by fed fish icv injected with either vehicle or NPY (N=6; **p<0.01, ***p<0.001). Data are represented as cumulative biting attempts in 15 min bins over a 1-hour period and were compared using Two-way ANOVA, with Bonferroni’s post-hoc analysis. **(C,D)** Representative micrographs of Dm region from the transverse sections across telencephalon showing p-ERK immunoreactive (p-ERK-ir) cells in fed fish icv injected with either vehicle or NPY **(E)** The number of p-ERK-ir cells in Dm of fed fish icv injected with either vehicle or NPY. The data were compared using unpaired t-test with Welch’s correction (N=6; **p< 0.01). **(F,G)** representative micrographs of one lobe of Dm region from the transverse sections across telencephalon showing phosphoNR1 (pNR1) immunoreactivity in starved fish icv injected with either glucose or co-injected with NPY along with glucose **(H)** Quantification of mean pNR1 intensity per neuron (au: arbitrary units) in Dm in starved fish icv injected with either glucose or co-injected with NPY along with glucose. The data were compared using unpaired t-test with Welch’s correction (12253 cells in vehicle+ glucose group and 11901 in NPY + glucose group; N=3; *** p< 0.001) [Scale bar 50 µm] **(I)** The traces represent relative change in fluorescence intensity (ΔF/F0) over time in response to 75 µM glutamate. The dark line represents mean ± SEM with the dotted lines denoting the limits for standard error. The coloured rectangular box indicates the duration for which glutamate was presented. The response of Dm neurons in starved fish ICV injected glucose (blue trace) and co-injected with NPY along with glucose (brown trace). **(J)** The max (ΔF/F0) as well as **(K)** AUC of response of Dm neurons in starved fish icv injected glucose compared with the starved fish co-injected with NPY along with glucose. The data were analysed using t-test with Welch’s correction. (N=6 telencephalons per group, *p<0.05).

To evaluate if NPY could increase the feeding drive, we intracerebroventricularly introduced NPY peptide into recently fed fish. As expected, vehicle-injected fed fish made infrequent biting attempts (338.8±70.4) while NPY injected animals showed increased feeding drive (983.7±75.84) that was comparable with starved animals (Fig. 7 B). However, there was no change in locomotion or anxiety-like behaviours in NPY-treated animals (Fig. S2).

### NPY signalling converges on the Dm neurons to decrease neuronal activity

Increased expression of NPY in response to starvation has been reported in several species (Loh et al., 2015), as also observed in the hypothalamic nuclei of zebrafish (Yokobori et al., 2012). Compared to fed zebrafish, the number of NPY positive neurons increased in the EN and the hypothalamic nucleus recessus lateralis (NRL) of starved animals (Fig. S6 A-F).

In transverse slices of the telencephalon, NPY-expressing EN soma was found to project to the Dm (Fig S6 G) similar to what was seen for CART-positive EN neurons. EN to Dm projections is consistent with previous reports (Turner et al., 2016). Together with DiI tracing experiments reported earlier (Fig. S4 B-D), the connectivity of periventricular hypothalamic nuclei and the EN to Dm (Fig. S6 H) raises the possibility of NPY signalling also converging on the Dm and opposing CART activity. We tested this possibility by evaluating the activity of the Dm neurons in response to NPY.

Icv administration of NPY to recently fed animals reduced the number of p-ERK positive Dm neurons (109.5±4.74) compared to vehicle controls (353.5±49.73; Fig. 7 C-E). Thus, the difference in the activity of Dm neurons observed between fed and starved animals (Fig. 2 D-F) can be partially explained by the opposing orexic and anorexic signalling by NPY and CART neuropeptides, respectively.

As CART signalling increased the phosphorylation of the NR1 subunit of NMDAR to increase neuronal activity, we tested if the reduction of Dm activity following NPY-treatment was a result of decreased NR1 phosphorylation. Indeed, glucose-stimulated (satiety mimic) pNR1 fluorescence (944.3±2.98) was substantially decreased (478.2±2.03) when NPY peptide was co-injected with glucose into the brain of starved fish (Fig. 7 F-H).

These data identify NPY-signalling, originating from the energy state sensitive populations of the hypothalamus and EN under energy poor metabolic states, in modulating zebrafish feeding behaviour. NPY-signalling converges on the Dm neurons, acts antagonistically to CART to reduce NR1 phosphorylation and decreases neuronal activity at the Dm.

### NPY signalling reduced the glutamate responsiveness of Dm neurons

To directly test if NPY signalling reduced the response of Dm neurons to glutamatergic stimulation, we evaluated the Dm activity in the Tg(NeuroD:GCaMP6f) zebrafish line. *Ex vivo* preparations of the intact telencephalon were dissected from starved animals treated earlier with either glucose (satiety mimic) or glucose and NPY via the icv route.

As expected, Dm of glucose injected animals responded to glutamate stimulation (maximum amplitude of response: 0.39±0.09 and magnitude of response: 75.45±12.93). However, the response to glutamate was substantially damped in the Dm of fish co-injected with glucose and NPY (maximum amplitude of response: 0.13±0.06 and magnitude of response: 28.43±4.7; Fig. 7 I-K).

These data show that NPY signalling alters the responsiveness of Dm neurons to glutamate and this altered activity state of Dm may be a neuronal representation of the energy deficit conditions. The orexic NPY signalling and the anorexic activities of the CART neuropeptide converge on the Dm neurons. Dm thus appears to be an important integrative centre whose activity is correlated with the homoeostatic regulation of feeding.

### Orexic signalling by NPY is mediated by the activation of calcineurin and inhibition of adenylyl cyclase

Our data identify PKA and PKC mediated phosphorylation of NR1 in the Dm neurons in response to CART in inducing satiety. Further, NPY signalling appears to oppose CART function, including the dephosphorylation of NR1 (Fig 7 H). Thus, we investigated NPY-induced signalling that could oppose CART function.

In mammals, NPY increases the activity of the protein phosphatase calcineurin (PP2B) (Chen et al., 2005; Sajdyk et al., 2008) and calcineurin is known to dephosphorylate NR1 (Choe et al., 2005). We blocked calcineurin activity using FK506 (calcineurin inhibitor) in NPY peptide-treated fed fish and evaluated their feeding behaviour. FK506 alone (222±57.39) did not show any significant effect on the number of biting attempts as compared to vehicle controls (291.4±79.85; Fig 8 A). However, the co-injection of FK506 along with NPY (368±50.97) reversed NPY induced orexia (954.7±67.55; Fig 8 A).

**Figure 8.**
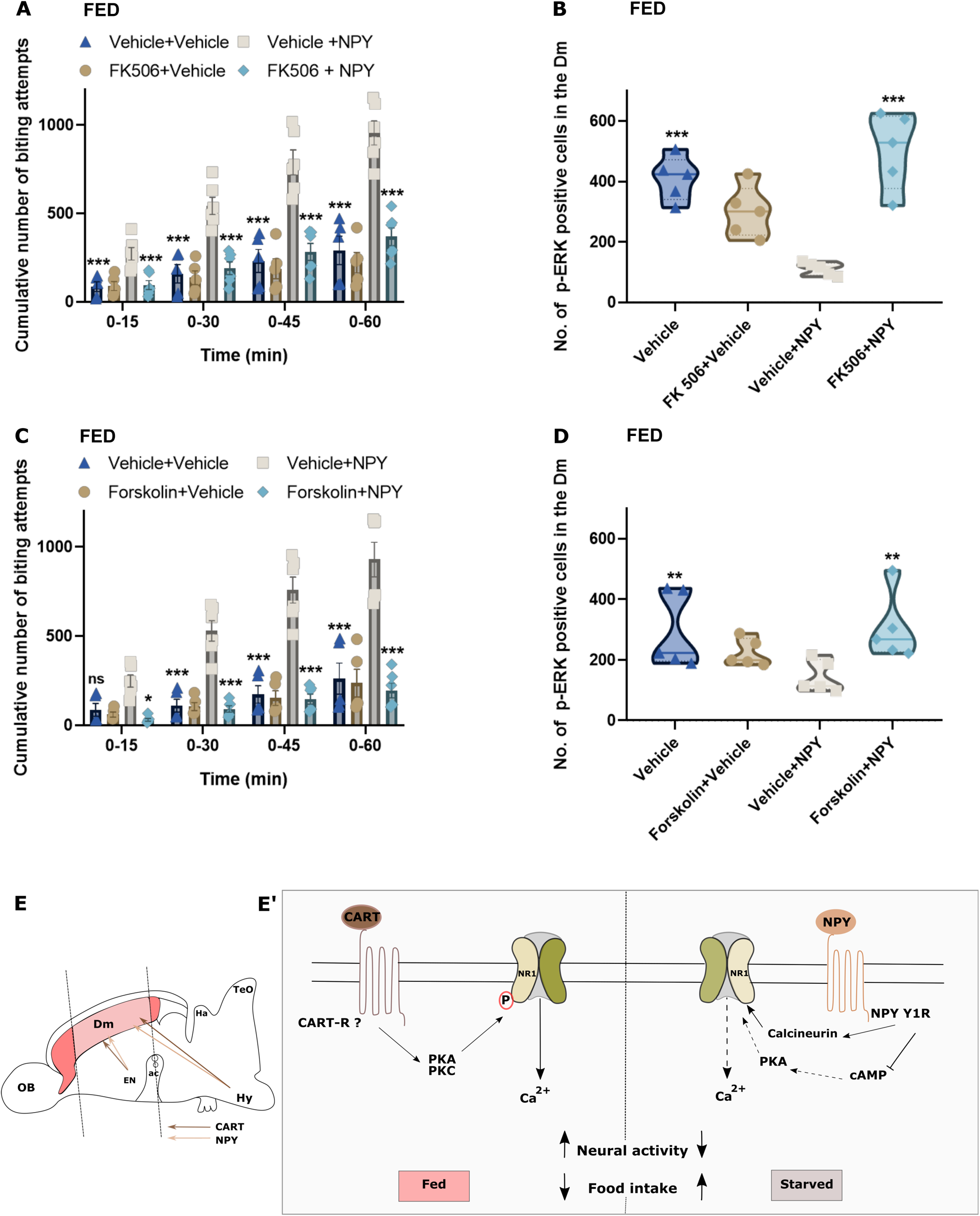
NPY inhibits adenylyl cyclase and activates of calcineurin to promote food intake. **(A)** The cumulative number of biting attempts made by fed fish icv injected with either vehicle, NPY, FK506 or co-injection of FK506 along with NPY. (N=5 animals per group). Data are represented as cumulative biting attempts in 15 min bins over a 1-hour period and compared using Two-way ANOVA, with Bonferroni’s post-hoc analysis for significance in comparison with the NPY treated fed fish(error bars represent ± SEM; *** p<0.001). **(B)** The number of p-ERK-ir cells in fed fish icv injected with either vehicle, NPY, FK506 or co-injection of FK506 along with NPY. Data were compared using Two-way ANOVA, with Bonferroni’s post-hoc analysis for significance in comparison with the NPY treated fed fish (N=5; *** p< 0.001). **(C)** The cumulative number of biting attempts made by fed fishicv injected with either vehicle, NPY, Forskolin or co-injection of Forskolin along with NPY (N=5 animals per group). Data are represented as cumulative biting attempts in 15 min bins over a 1-hour period and were compared using Two-way ANOVA, with Bonferroni’s post-hoc analysis for significance in comparison with the NPY treated fed fish (error bars represent ± SEM; *** p<0.001, *p<0.05). **(D)** The number of p-ERK-ir cells in fed fish icv injected with either vehicle, NPY, Forskolin or coinjection of Forskolin along with NPY. Data are compared using Two-way ANOVA, with Bonferroni’s post-hoc analysis for significance in comparison with the NPY treated fed fish(N=5; ** p< 0.01). **(E)** Schematic showing CART and NPY projections to Dm from interoceptive regions in zebrafish brain. [OB: olfactory bulb; EN: Entopeduncular nucleus; ac: anterior commissure; Ha: Habenula; Hy: Hypothalamus; TeO: Optic tectum] **(E’)** Summary of modulatory changes induced by CART and NPY leading to a corresponding change in neural activity and food intake under fed and starved conditions, respectively. Under energy enriched conditions (Fed), CART signalling at the Dm is increased. CART activates PKA and PKC to increase the phosphorylation of NR1 subunit of NMDARs and thereby increases NMDAR activity in Dm neurons. CART mediated increase in Dm neuronal activity is correlated with a decrease in food intake in zebrafish. In contrast, under energy deprived conditions, NPY signalling at the Dm is increased. NPY acting via Y1R reduces PKA activity and activates calcineurin to decrease phosphorylation of NR1, in turn, decreasing NMDAR activity. NPY mediated reduction in Dm neuronal activity is correlated with increased food intake in zebrafish.

To test if blocking calcineurin activity affects NPY induced reduction in Dm activity, we co-injected FK506 and NPY peptide in fed fish. Treatment with the FK506 alone (299.8±38.11) did not affect the number of p-ERK positive neurons in the Dm of fed fish (409.6±32.62; Fig 8 B). However, blocking calcineurin activity in NPY treated fish (502.4±56.7) abrogated NPY-induced (111±9.38) reduction in p-ERK positive neurons in Dm.

We have demonstrated that NPY function in regulating food intake in zebrafish is via the Y1 receptor (Fig. 7 A). NPY Y1R is G protein receptor that inhibits adenylyl cyclase activity (Molosh et al., 2013). A fall in cAMP production by adenylyl cyclase is expected to reduce the activation of PKA, and PKA is critical for CART signalling (Fig 4 A,B). Therefore, we activated adenylyl cyclase with Forskolin (an adenylyl cyclase agonist) in fed fish co-treated with NPY. Forskolin alone (236±77.40) did not influence the number of biting attempts as compared to vehicle controls (261.8±86.98; Fig. 8 C); however, application of Forskolin in NPY co-injected fed fish reversed (194.7±40.7) NPY induced orexia (929±96.86; Fig. 8 C).

Forskolin co-administered along with NPY in fed fish, via the icv route, alleviated (304.1±49.90) NPY-mediated reduction of p-ERK positive neurons in the Dm of fed animals (145.1±23.68; Fig. 8 D). Treatment with forskolin alone (223.8±20.05) was comparable to vehicle-injected fed fish (296.6±56.19; Fig 8 D).

Collectively, these experiments demonstrate that orexic NPY signalling is mediated by the downregulation of cAMP activity and upregulation of calcineurin activity, which decreases the activity of Dm neurons via reduced phosphorylation of NR1. NPY-signalling is antagonistic to CART-induced phosphorylation of NR1 and the energy-state dependent expression of CART/NPY results in the bimodal regulation of Dm activity and feeding behaviour.

## DISCUSSION

The flexibility of innate, motivated behaviours is driven by context-dependent recruitment and outputs of the underlying neural circuits (Sternson, 2013; Flavell et al., 2020). Neuromodulator-based signalling has emerged as a key mechanism facilitating reconfiguration of circuit activity (Lee and Dan, 2012). For instance, state changes between component behaviours in *C. elegans* foraging is regulated by the differential recruitment of circuits by the opposing activities of serotonin and the PDF neuropeptide (Flavell et al., 2013). Neuropeptide activities may also change the response of specific neurons or circuits. Peptidergic modulation of the worm ASH neuron mediates feeding state-dependent nociceptive responses (Ezcurra et al., 2016). Similarly, NPY-signalling regulates the sensitivity of zebrafish olfactory receptor neurons in response to hunger (Kaniganti et al., 2021).

Homoeostasis necessitates altering food intake in response to changing energy stores. Therefore, the neural circuits regulating food intake need to be dynamic and responsive to physiological states. Neuropeptidergic signalling may represent energy state information and facilitate biochemical modulation of synaptic transmission to regulate food intake-associated behaviours. Indeed, a diverse and complicated network of peptidergic activity, both peripheral and central, influence the plasticity of neural circuits regulating food intake (Crespo et al., 2014). However, the neuropeptide-induced signalling facilitating the integration of opposing energy state signals to generate appropriate behavioural outputs remains poorly understood.

In the mammalian Arc, the POMC/CART-expressing neurons of the Arc release α-MSH onto the MC4R-expressing neurons of the paraventricular hypothalamic nucleus (PVH) to enhance glutamatergic neurotransmission and induce satiety (Fenselau et al., 2017). Conversely, the GABAergic AgRP/NPY neurons on the other hand provide an inhibitory tone onto MC4R-expressing PVH neurons to promote food intake (Fenselau et al., 2017). While opposing inputs from the Arc converge onto the PVH neurons, the intracellular mechanisms recruited by these peptidergic neuromodulators are inadequately characterised.

We demonstrate the convergence of antagonistic intracellular signalling mediated by the anorexic neuropeptide CART and the orexic NPY on a zebrafish telencephalic neuronal population to reconfigure its activity. Energy state sensitive expression of the neuropeptides impose opposing biochemical constraints on their target neurons at the dorsal telencephalon via the modulation of NMDAR function and regulate the feeding behaviour.

We show that not only do the endogenous activities of CART and NPY modulate the feeding drive, but the intracerebroventricular delivery of these peptides acutely alters food intake in the expected direction. We limit our analysis to intracerebroventricular (icv) delivery of bioactive molecules and pharmacological agents to focus on central activities and avoid potential confounds originating from peripheral inputs. Consistent with earlier work (Subhedar et al., 2011; Mukherjee et al., 2012; Yokobori et al., 2012; Akash et al., 2014), we also find that expression of CART peptide increases, under energy-rich conditions, in energy state interoceptive regions of the periventricular hypothalamus and the EN of zebrafish while NPY expression increases upon starvation.

Anatomical analysis indicated that the peptidergic neurons of the periventricular hypothalamus and the EN project to the Dorsomedial telencephalon (Dm) and suggested that the latter population may be the target for CART or NPY, under different metabolic states. The connectivity observed in our tracing studies is in line with observations reported in other teleosts (Kanwal et al., 1988), including zebrafish (Turner et al., 2016; Yáñez et al., 2017). Thus, our subsequent studies focussed on the neuronal activity at the Dm, in response to CART and NPY.

Using p-ERK as a readout of recent neuronal activity and calcium imaging of a genetically encoded calcium sensor, we found that energy-rich states resulting from recent feeding (and consequent increase in endogenous CART neuropeptide) or exogenous application of CART peptide to the brain led to heightened activity of the Dm neurons. The increased activity of the Dm neurons was found to be NMDAR dependent and blocking NMDAR signalling reduced Dm activity and CART-mediated anorexia.

Studies in rodents have previously reported the involvement of NMDA receptors in the regulation of food intake. Under satiety conditions, NMDA receptors in the hindbrain are required for CCK induced reduction of food intake (Wright et al., 2011). NMDAR function has also been implicated in leptin (Neyens et al., 2020) and PACAP (Resch et al., 2014) mediated reduction in food intake. Furthermore, NMDAR activity in the lateral hypothalamus regulates food intake in hungry animals (Stanley et al., 2011), while it is necessary for the activity of the AgRP-expressing neurons to establish the starvation state (Liu et al., 2012). In addition, CART function in processing innate fear in the central amygdala (Rale et al., 2017) and nociception by sensory neurons (Chiu et al., 2010) also involve NMDAR function and may represent a common modality in CART signalling.

Further investigations identified PKA and PKC activities in mediating CART induced anorexia since inhibiting the activity of either kinases led to the restoration of food intake and decreased neuronal activation in the Dm. Using phosphospecific NR1 antibodies we found that CART induced activation of PKA and PKC results in phosphorylation of NR1 on specific residues.

Phosphorylation of NR1 by PKC and PKA has been known to enhance NMDAR signalling by increasing both the Ca^2+^ permeability (Skeberdis et al., 2006) and the opening probability of the receptor complex (Lan et al., 2001). Thus CART-induced phosphorylation is likely to increase the responsiveness of Dm neurons to glutamatergic inputs and is consistent with our results showing heightened calcium response of Dm neurons upon stimulation by glutamate.

The modulation of Dm activity by endogenous CART release in response to energy-rich conditions was demonstrated using calcium imaging of *ex vivo* whole-brain preparations where the inputs from the hypothalamus and EN are preserved. Brains isolated from starved animals (low endogenous CART and high endogenous NPY) failed to respond to glutamate when maintained under glucose deprived conditions. However, incubation with glucose restored glutamate responsiveness and suggests altered inputs (including CART) from the glucose-sensing interoceptive areas. While this response may involve modulatory inputs to the Dm not limited to CART, it is consistent with our observations of CART function. Importantly, as the experimental preparation precludes peripheral signals of energy state, the modulation of Dm activity appears to be primarily driven by neuronal inputs.

NPY containing neurons of the EN and periventricular hypothalamus also project to the Dm. Thus, the Dm neurons could function as a common substrate for orexic and anorexic inputs and their activity could be dynamically tuned to the prevailing energy states. Indeed, both starvation and exogenous application of NPY to fed fish (low levels of endogenous NPY) led to a reduction in Dm activity with a concomitant increase in the feeding drive.

Our investigation reveals that NPY opposes CART-mediated elevation of NR1 phosphorylation and consequent increase in Dm activity and anorexia by reducing cAMP production (thereby reducing PKA activation) and activating the phosphatase calcineurin. Our observations are consistent with NPY-mediated activation of calcineurin (Chen et al., 2005; Sajdyk et al., 2008) and calcineurin mediated dephosphorylation of NR1(Choe et al., 2005). Further, NPY Y1R mediated reduction in cAMP and PKA has been shown to decrease NMDA-mediated postsynaptic currents (Molosh et al., 2013) in the neurons of the basolateral amygdala.

Our data suggest that Dm neurons are modulated by the opposing activities of CART and NPY to achieve two distinct activity states (Fig. 8 E). Dm neurons, in zebrafish, may represent “second order” neurons, immediately downstream of the interoceptive, “first order” population in the hypothalamus and EN. While our experiments are unable to distinguish whether the same individual neurons respond differentially to CART and NPY, the tight correlation of Dm activity with alterations in food intake behaviour suggest a pivotal role for Dm neurons in regulating energy homoeostasis (Fig. 8 E). A major limitation in delineating CART function has been the lack of well characterised receptors and selective pharmacology. This drawback has limited our analysis of Dm neurons. However, collateral evidence from the gustatory systems underscores the role of Dm in food intake. In a range of teleosts, the preglomerular tertiary gustatory nucleus is reciprocally connected to the Dm (Kanwal et al., 1988; Kato et al., 2012). Though the downstream targets of the Dm relevant to the feeding drive are unknown, it appears CART/NPY activities modulate the responsiveness of the Dm to glutamatergic inputs to release or inhibit food intake.

The Dm is considered to be homologous to the mammalian amygdaloid complex (Wullimann and Mueller, 2004; Porter and Mueller, 2020). Several recent reports identify the mammalian amygdalar complex in the regulation of feeding behaviours (Padilla et al., 2016; Kim et al., 2017; Izadi and Radahmadi, 2021). Consistent with our findings, the amygdala receives projections from both the NPY and CART neurons of the hypothalamic Arc (Hwang and Lee, 2018) and employs NPY Y1 mediated signalling to increase the food intake (Padilla et al., 2016). Additionally, projections from dopamine receptor-positive neurons in prefrontal cortex are known to activate medial BLA to increase food intake (Land et al., 2014). The amygdala is also involved in hindbrain satiety circuits, where PKC-δ+ neurons in the central amygdala (CeA) integrate multiple anorexigenic signals to inhibit food intake (Cai et al., 2014). Conversely, Htr2a+ neurons of the CeA have been shown to stimulate food intake (Douglass et al., 2017). In addition, several anorexigenic neuromodulators, when administered to the CeA have been shown to affect food intake (Fekete et al., 2007; Beckman et al., 2009; Kovács et al., 2012). A recent study has identified appetitive information specific projections from the BLA to the CeA and isolated specific CeA subpopulations that differentially mediate fear processing and feeding-associated behaviours (Kim et al., 2017).

While the microcircuitry and the diversity of neuronal subtypes in the zebrafish Dm remain unknown, the feeding regulation-associated CART/NPY-responsive neurons could be distinct from those processing emotions. Consistent with this, no difference in anxiety-like behaviours was seen in fish under fed and starved states or treated with either of the neuropeptides (Fig. S2). The CART/NPY-responsive neurons are a subpopulation restricted to a specific region of the Dm - starting at the level of the anterior commissure and extending 250 µm rostrally (Fig. 2 G-I). Identification of neuronal subtypes within the Dm and development of intersectional genetics will be critical in characterising the CART/NPY-responsive microcircuits in the future.

This study investigates neural mechanisms mediating the opposing activities of the neuropeptides CART and NPY in regulating the feeding drive in zebrafish and identifies plasticity in a group of neurons in the zebrafish forebrain. CART and NPY inputs converge on Dm neurons to antagonistically modulate NMDAR function, resulting in an energy state-dependent alteration in glutamatergic neurotransmission that is tightly correlated with the behavioural switch in food intake.

## ETHICS APPROVAL

All protocols used in this study were approved by Institutional Animal Ethics Committee of IISER Pune.

## AVAILABILITY OF DATA AND MATERIAL

All data generated or analysed during this study are included in this published article. The raw data are available from the corresponding author on reasonable request.

## AUTHOR CONTRIBUTIONS

Conceptualization: D.B., A.M., N.S. and A.G.; Investigation and formal analysis: D.B., A.M. and T.K.; Generation of preliminary data: D.W. and A.B.; Writing – original draft: D.B., A.M. and A.G.; Writing – review and editing: D.B., A.M., N.S. and A.G.; Funding Acquisition: A.G.. All authors gave final approval for publication and agree to be held accountable for the work performed therein.

## ACKNOWLEDGMENTS

The authors thank Dr. G. Deshpande, Princeton University, Dr. S. Rath, IISER Pune and Dr. R. Rajan, IISER Pune for their critical reading of this manuscript and discussions. The Tg(NeuroD:GCaMP6f) line was a gift from Dr. V. Thirumalai, NCBS, Bengaluru. N. Tiwari, IISER Pune is acknowledged for contributing to the development of the feeding assay. The authors acknowledge the IISER Pune Microscopy Facility and the National Facility for Gene Function in Health and Disease (NFGFHD) at IISER Pune for access to equipment and infrastructure.

