## Supplemental Information for "Convergent energy state-dependent antagonistic signalling by CART and NPY modulates the plasticity of forebrain neurons to regulate feeding in zebrafish"

**This file includes:**

Figures S1 to S6

Supplementary Materials and Methods

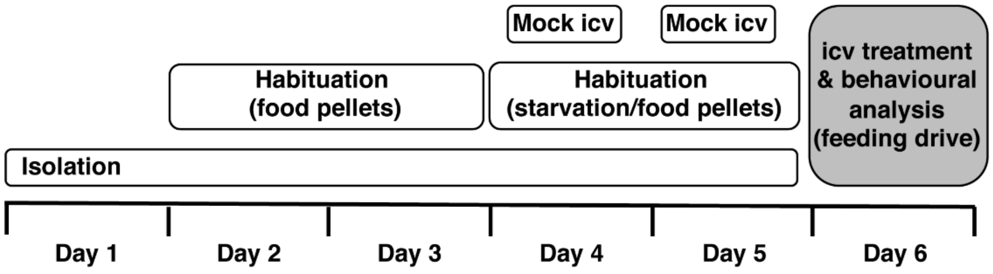


**Figure S1: Experimental design for the analysis of feeding drive**

Schematic of experimental design and timeline for feeding behavioural experiment. Details of the protocol, pharmacological and bioactive agents, and doses used are provided in Methods section.

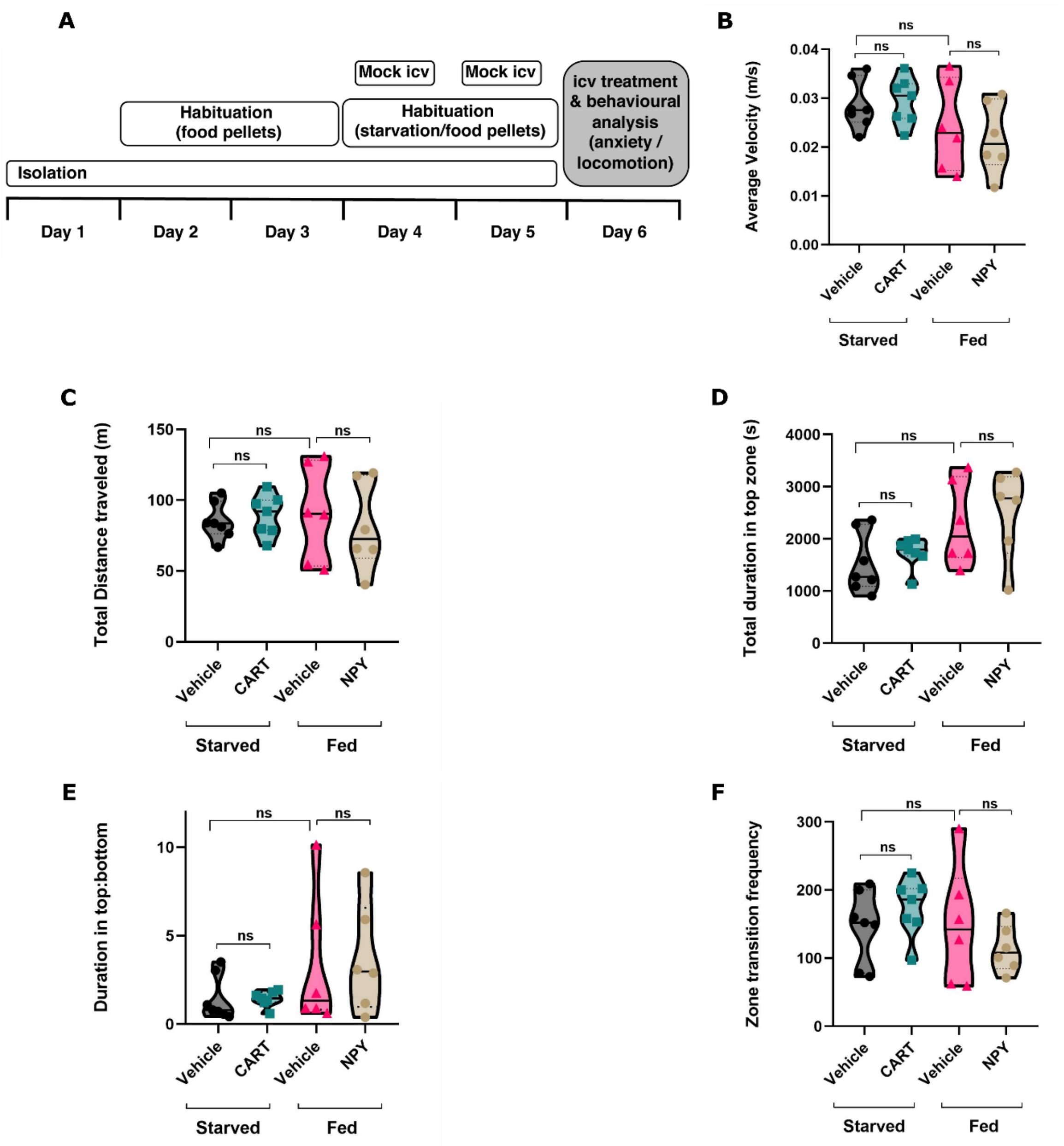


**Figure S2: Analysis of anxiety-like behaviours and locomotion kinematics**

**(A)** Schematic of experimental design and timeline used for the evaluation of anxiety-like behaviours and locomotion. **(B)** Average velocity, **(C)** total distance travelled, **(D)** duration in the top zone, **(E)** duration in top:bottom, **(F)** transition frequency between the top and botton zones were assessed under different physiological energy states or upon icv administration of CART and NPY peptides. The data were compared using unpaired, t-test with Welch's correction (N=6 animals in each group; ns, not significant). Details of the protocol, pharmacological and bioactive agents, and doses used are provided in Supplementary Methods section.

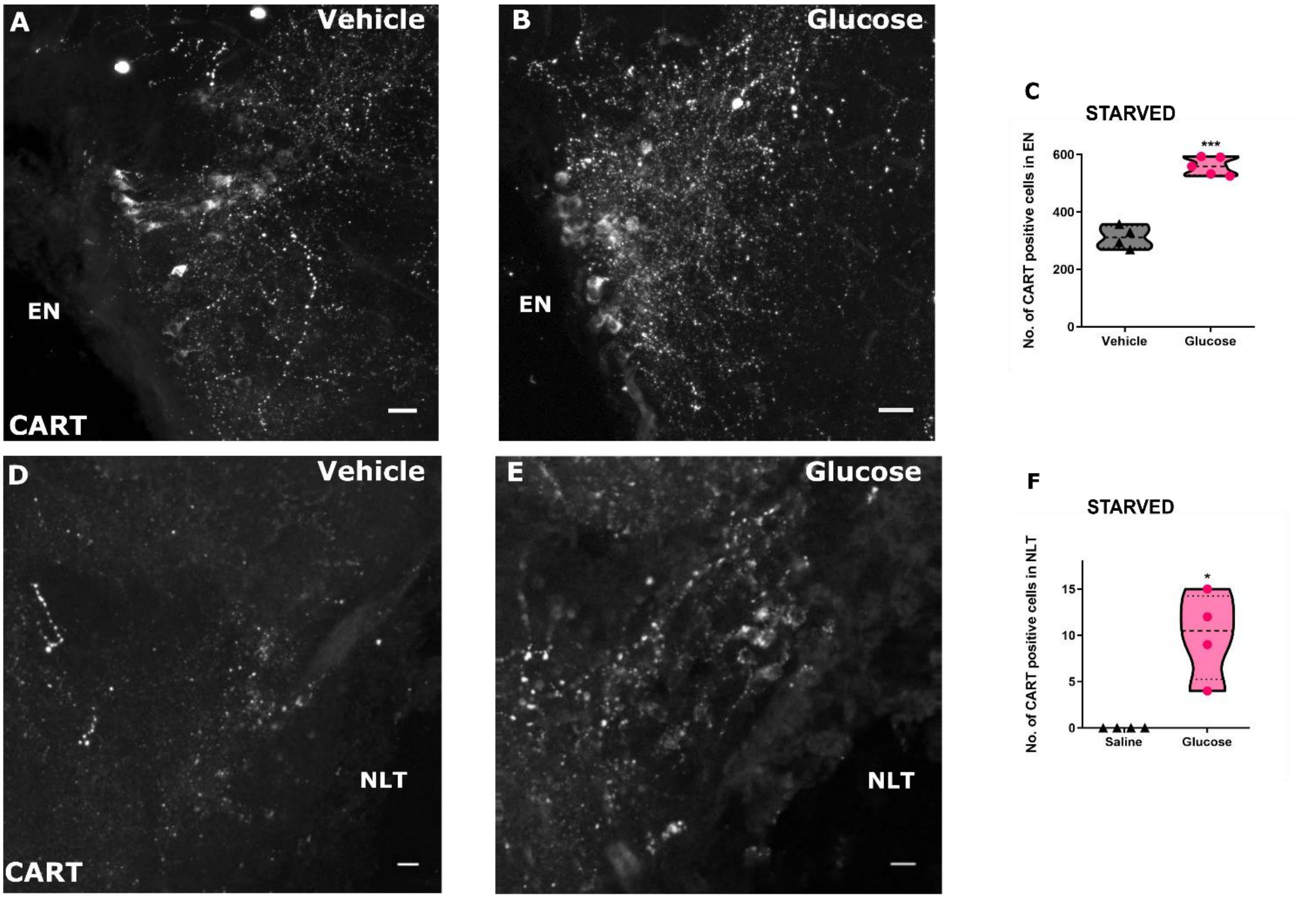


**Figure S3: CART expression is upregulated in the EN and the NLT upon icv glucose administration**

Representative micrographs of the entopeduncular nucleus (EN) **(A,B)** and the nucleus lateralis tuberis (NLT) **(D,E)** from transverse sections of the zebrafish brain showing CART immunoreactive cells in starved fish receiving icv injection of either vehicle or glucose. The quantification of the number of CART positive cells in EN **(C)** and NLT **(F)** in starved fish icv injected with either vehicle or glucose. The data were compared using unpaired t-test with Welch's correction (N=4/5; * $p< 0.05$; *** $p< 0.001$)

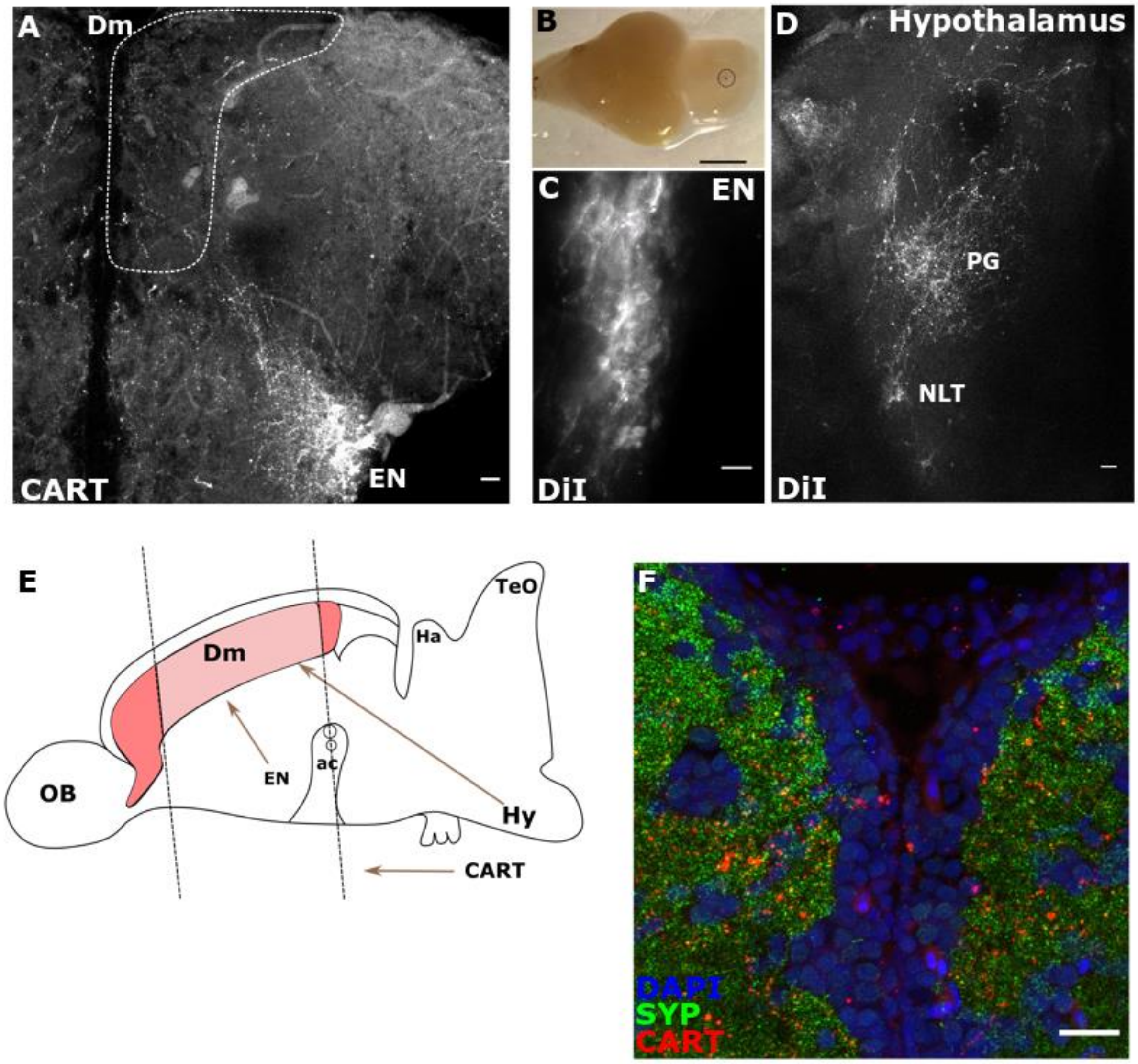


**Figure S4: Dm receives inputs from the EN and the periventricular hypothalamus**

**(A)** Transverse section of telencephalon showing CART immunoreactive neurons in the entopeduncular nucleus (EN) and CART-containing fibre tracts projecting to Dm region (marked by dotted line). **(B-D)** Analysis of Dm connectivity by DiI labelling **(B)** Site of DiI application in the Dm (dotted circle). **(C,D)** Representative photomicrographs of the optical sections showing DiI labelling in EN and nucleus lateralis tuberis (NLT) neurons. **(E)** Schematic showing the connectivity of EN and the periventricular hypothalamic projections to the Dm (shaded region) [OB, olfactory bulb; EN, entopeduncular nucleus; ac, anterior commissure; Ha, habenula; Hy, hypothalamus; TeO, optic tectum] **(F)** Presynaptic localisation of CART in the Dm. Representative micrographs of Dm showing colocalization of synaptophysin (SYP; green) and CART (red) along with nuclei labelled with DAPI (blue). Scale bar: 1000 µm in B; 25 µm in A; 20 µm in C,D,F.

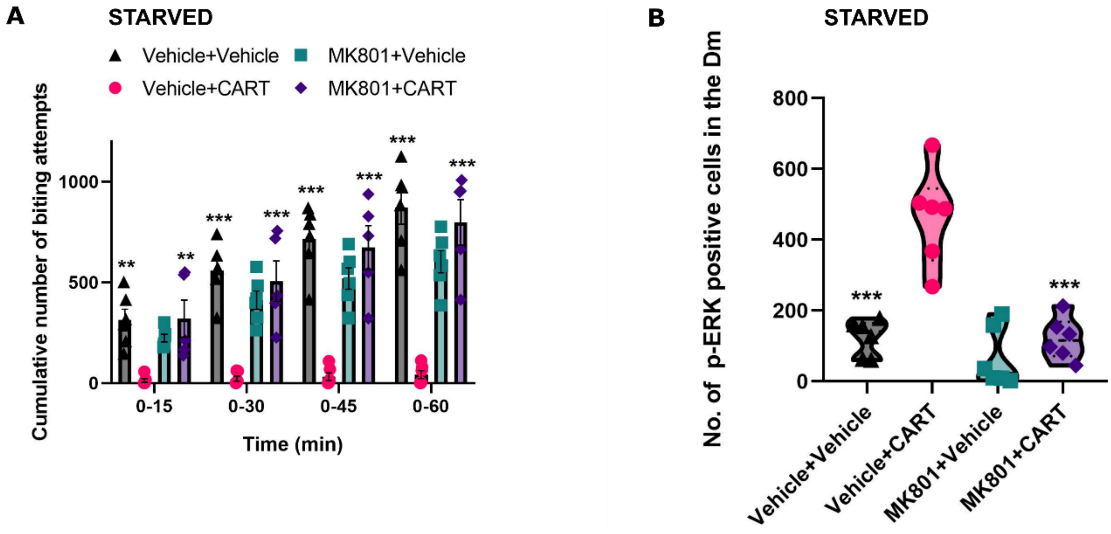


**Figure S5: CART-induced anorexia and activation of Dm neurons require NMDAR signalling**

**(A)** The cumulative number of biting attempts made by starved fish icv injected with either vehicle, CART peptide, MK801 or coinjected with MK801 and CART peptide. Data are represented as cumulative biting attempts in 15 min bins over a 1-hour period and compared using two-way ANOVA, with Bonferroni's post-hoc analysis analysis for significance in comparison with the CART treated starved fish (error bars represent ± SEM; N=5 animals for MK801+CART and N=6 animals for the other groups; $^{**}p<0.01$, $^{***}p<0.001$). **(B)** The number of p-ERK immunoreactive cells in starved fish icv injected with either vehicle, CART peptide, MK801 or coinjected with MK801 and CART peptide. The data were compared using two-way ANOVA, with Bonferroni's post-hoc analysis analysis for significance in comparison with the CART treated starved fish (N=6 animals per group; $^{***}p< 0.001$).

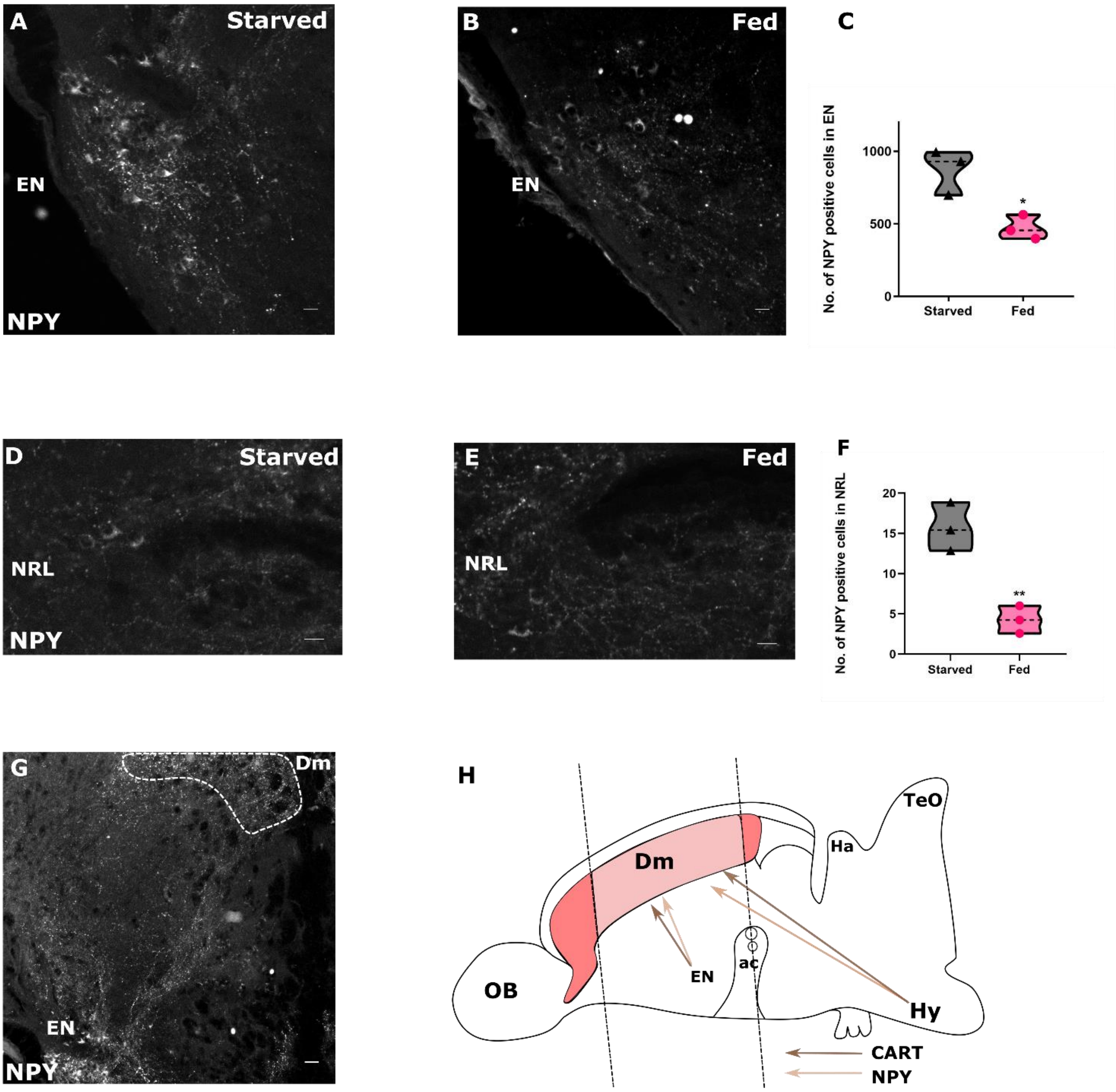


**Figure S6: NPY expression is upregulated in the EN and the NRL in response to starvation.**

Representative micrographs of the entopeduncular nucleus (EN) **(A, B)** and the nucleus recessus lateralis (NRL) **(D, E)** regions from transverse sections of the zebrafish brain showing NPY immunoreactive cells in starved and fed fish. The quantification of number of NPY positive cells in EN **(C)** and NRL **(F)** in starved and fed fish. The data were compared using unpaired t-test with Welch's correction (N=3 animals per group; ** $p < 0.01$). **(G)** Transverse section of telencephalon showing NPY immunoreactive neurons in EN and NPY-containing fibre tracts projecting to

Dm region (marked by dotted line). **(H)** Schematic showing connectivity of EN and Hypothalamic CART and NPY projections to the Dm (shaded region) [OB, olfactory bulb; EN, entopeduncular nucleus; ac, anterior commissure; Ha, habenula; Hy, hypothalamus; TeO, optic tectum]**.** Scale bar: 25 µm.

### SUPPLEMENTARY MATERIALS AND METHODS

#### Analysis of locomotion and anxiety-like behaviours

Fish were isolated from home tanks and housed singly in the experimental tank. Fig. S2A outlines the protocol followed for all feeding behaviour experiments. Briefly, for first three days of habituation, the experimental tanks were moved to the recording chamber and the fish were allowed to feed on coloured food pellets (~ 15±5 pellets; Taiyo) for 1 hour before returning to the housing chamber. For habituation to injection and handling stress, the fish were anesthetised and received icv saline (0.9% NaCl) injection (day 4) or a mock injection (day 5). They were allowed to recover in recording chamber for 1 hour, before returning the tanks to the housing chamber. Fish that were food deprived for 2.5 days starting from day 3 are referred to as 'starved' while those that continued to receive food pellets are referred to as 'fed'. On the day of the experiment, fish were anesthetised, icv injected with appropriate reagents/vehicle and returned to experimental tanks. Following a recovery period of 15 mins, the behaviour of the fish (taken one at a time) was monitored in the absence of food pellets using a video recorder (Sony Handycam) for one hour. The videos were then analysed using Ethovision-XT software (Noldus) for parameters indicating anxiety-like behaviour (i.e total duration in top half of the tank, ratio of time spent in top vs bottom half of the tank and frequency of transitions to the bottom half) and locomotion status (viz. average swimming velocity and total distance travelled).

#### DiI Tracing

Adult zebrafish of either sex were anaesthetized, the brains were exposed, fixed in PFA (4%, pH 7.4) and post-fixed in the same solution. DiI crystals were placed in the dorso-medial telencephalon (Dm). The brains were kept in paraformaldehyde solution at room temperature in dark for 7 days and were processed for sectioning or for whole brain imaging using confocal microscope (Zeiss LSM 710).

#### Immunofluorescence

Fish were subjected to icv injection with glucose (40 nmol in 1x PBS) or vehicle and allowed to recover for 30 mins. After recovery, fish were anesthetised and craniotomised to expose the dorsal surface of the brain. Samples were fixed in 4% PFA overnight at 4 $^{o}$C. For experiments comparing NPY levels, well-fed fish or fish starved for 3 days were anesthetised and craniotomised prior to fixation in Bouin's fixative (71.4% saturated solution of picric acid, 23.8 % formalin & 4.7% glacial acetic acid). After fixation for 12-14hrs brain was dissected and cryoprotected with 25% sucrose prior to sectioning. Serial 10 µm thick sections of the entire telencephalon were mounted onto lysine coated slides and stored at -40 $^{o}$C till further processing. Sections were processed for staining with the following primary antibodies: anti-CART antibody (1:500, gift from Drs. Lars Thim and Jes Clausen, Novo Nordisk, Denmark), anti-Synaptophysin (1:100, Invitrogen, Cat. No. PA1-1043), anti- NPY antibody (1:5000, Sigma Aldrich, N9528). Details of the staining protocol are as given in the methods section of the main text.

Secondary antibodies were added and incubated at room temperature (RT) for 3 hrs in dark. The following secondary antibodies were used: Anti-rabbit Alexa Flour 488/568 (1:500, Invitrogen, Cat. No. A-11034/ A-11036) and anti-mouse Alexa Flour 488/568 (1:500, Invitrogen, Cat. No. A-11029/A-11031). Sections were washed three times and mounted in media (0.5% N propyl gallate, 70% glycerol, 1M Tris pH 8.0) containing 4′,6-diamidino-2-phenylindole (DAPI) (Invitrogen, Cat. No. D1306). Sections were observed under Axioimager Z1 (Zeiss) epifluorescence microscope and representative images were acquired using confocal microscope (SP8, Leica or LSM 780, Zeiss). In order to ensure reliable comparisons across different groups and maintain stringency in tissue preparation and staining conditions, all the brain sections were processed concurrently under identical conditions. Details of cell count analysis and statistical analysis can be found in methods section of the main text.
